# Mitotic spindle assembly is controlled by the APC/C localized at the centrosome

**DOI:** 10.1101/2020.03.23.003210

**Authors:** Thomas Tischer, Jing Yang, David Barford

## Abstract

The control of protein abundance is a fundamental regulatory mechanism during mitosis. The anaphase promoting complex/cyclosome (APC/C) is the main protein ubiquitin ligase responsible for the temporal regulation of mitotic progression. It has long been speculated that the APC/C might fulfil other functions including assembly of the mitotic spindle. Here, we show that the APC/C localizes to centrosomes, the organizers of the eukaryotic microtubule cytoskeleton, specifically during mitosis. The APC/C is recruited to spindle poles by the centrosomal protein Cep152, and we identified Cep152 as both a novel APC/C interaction partner, and as an APC/C substrate. Importantly, this revealed a mitotic function of Cep152 that is reciprocally regulated by the APC/C. A destruction-defective mutant of Cep152 showed that the timely regulation of Cep152 levels at the centrosome controls spindle assembly and chromosome segregation. The APC/C-mediated degradation of Cep152 at the centrosome releases Cep57 from an inhibitory complex to enable its interaction with pericentrin, a critical step in promoting microtubule nucleation. Thus, our study extends the function of the APC/C from being a regulator of mitosis to also acting as a positive governor of spindle assembly. The APC/C thereby integrates control of these two important processes in a temporal manner.

## Introduction

During mitosis a mother cell divides into two daughter cells that both inherit identical genetic material. This process needs to be tightly controlled to ensure that chromosomes are equally distributed. Mitosis is therefore regulated by multiple mechanisms that work together to coordinate a successful cell division. One important regulatory mechanism is ubiquitin-mediated protein degradation, which controls the abundance of different cell cycle regulators during mitosis. The anaphase promoting complex/cyclosome (APC/C) is an E3 ubiquitin ligase governing this process. APC/C activity is coordinated by the correct attachment of chromosomes to the mitotic spindle. During recent years the molecular structure and mechanisms of the APC/C have been defined (Alfieri et al., 2017; Watson et al., 2019), however relatively little is known about the role of its intracellular localization. The APC/C has been observed at kinetochores during an active spindle assembly checkpoint (SAC) (Acquaviva et al., 2004) and at chromosomes (Sivakumar et al., 2014). Additionally, studies suggest that localized protein degradation plays an important role during mitosis. The APC/C substrate cyclin B, for example, was reported to be degraded first at the spindle poles (Clute and Pines, 1999; Huang and Raff, 1999). Consequently, the APC/C was shown to localize to spindle poles with this recruitment being dependent on the protein NuMA, and the motor protein dynein (Ban et al., 2007; Tugendreich et al., 1995). Recently, it has also been demonstrated that the APC/C co-activator Cdh1 is recruited to centrosomes by the centrosomal protein Cep192 in *Drosophila* (Meghini et al., 2016). It has therefore been speculated that the APC/C could be directly involved in spindle assembly.

Centrosomes are cellular organelles and the major organizers of the microtubule cytoskeleton in eukaryotic cells (Gonczy, 2012). In higher eukaryotes, centrosomes consist of two cylindrical shaped structures named centrioles that are embedded in a protein-rich matrix, the pericentriolar material (PCM). Centrosomes are duplicated in S-phase when the two tethered mother centrioles each gives rise to exactly one new daughter procentriole. In mitosis, the segregated centrosomes form the centre of two spindle poles that organize the mitotic spindle apparatus (Varmark, 2004). At anaphase, both newly formed daughter cells inherit one centrosome, and the centrosome’s centriole and procentriole split apart in a process called disengagement. These subsequently serve as new mother centrioles during the next centrosome duplication cycle in the following S-phase (Nigg and Holland, 2018; Stearns, 2001). Centrosomes dynamically recruit additional proteins around them to form the PCM, including pericentrin (PCNT), Cdk5rap2/Cep215 and gamma-tubulin (γTub), that are required for microtubule nucleation (Gupta and Pelletier, 2017; Kim et al., 2019; Kim and Rhee, 2014; Thawani et al., 2018; Wieczorek et al., 2020) and, consequently, for mitotic spindle formation. The PCM increases in size before mitosis (Lawo et al., 2012; Sonnen et al., 2012; Watanabe et al., 2019; Woodruff et al., 2014).

In this study we investigate the localization and function of the APC/C at spindle poles and its role in spindle assembly. Using a combination of super-resolution and confocal microscopy, cell biology and biochemical assays, we show that during mitosis the APC/C is localized to the PCM, organized into a ring-like structure. We establish that the main interacting protein responsible for the localization of the APC/C to the centrosome is Cep152, which we additionally show to be a novel APC/C substrate. Cep152 is locally ubiquitinated by the APC/C at the centrosome during mitosis, which in turn decreases the localization of the APC/C itself at the centrosome. This negative feedback loop controlling APC/C localization is required for proper assembly of the mitotic spindle. Stabilization of a Cep152 mutant that cannot be targeted by the APC/C results in reduced microtubule nucleation and increased chromosome mis-segregation. This phenotype is caused by sequestering the PCNT-binding protein Cep57 into an inhibitory complex consisting of Cep57-Cep63-Cep152 (Lukinavicius et al., 2013). Once Cep152 auto-regulates its removal from the centrosome by the APC/C, Cep63 is removed from the centrosome as well and Cep57 is liberated, which can now recruit PCNT, aiding microtubule nucleation. Hence, our work shows for the first time a role of Cep152 during mitosis and extends the function of the APC/C from being a critical director of cell cycle progression, to additionally also functioning as a positive regulator of mitotic spindle assembly.

## Results

### The APC/C is a component of the mitotic centrosome

During interphase, the APC/C is localized throughout the cell, with an increased signal present around the centrosome (Supplementary Figure 1a). This localization resembles the pattern of centriolar satellites, small membrane-less granules that surround the centrosome, and which were proposed to function as storage for centrosomal proteins (Hori and Toda, 2017; Lopes et al., 2011; Prosser and Pelletier, 2020). Immunofluorescence of the essential satellite component, pericentriolar material 1, (PCM1) (Kubo et al., 1999), together with the APC/C subunit APC2, revealed that the APC/C colocalized with centriolar satellites (Supplementary Figure 1a). This is in agreement with a recent proteomics study of centriolar satellites that also detected the APC/C to be present in these structures (Gheiratmand et al., 2019). To allow for better visualization of the APC/C signal during mitosis, we pre-extracted soluble cytoplasmic proteins before fixation of the cells (Tugendreich et al., 1995). Using antibodies against APC2 and APC3 allowed us to specifically visualize the APC/C at the spindle poles (Figure 1a). Although pre-extraction reduces the cytoplasmic signal from the stained proteins, it can lead to apparent mis-localization of the remaining protein (Melan and Sluder, 1992). To verify our results, we therefore fixed cells without pre-extraction. This showed a similar localization for APC2 (Supplementary Figure 1b, c). Expression of eGFP-tagged APC2 or APC3 in HEK293 cells confirmed the localization results we obtained by antibody-mediated immunofluorescence (Supplementary Figure 1d). Free eGFP alone does not localize to centrosomes (Supplementary Figure 1d, lower panel). Additionally, we also stained human retina pigmented epithelium (RPE1) cells (Supplementary Figure 1c) and used another APC2 antibody (“APC2 abcam”) which recognizes a different epitope for staining. Both experiments confirmed our localization observations using HEK293 cells (Supplementary Figure 1b). Using an siRNA against APC2, and probing for the APC2 protein by immunoblotting and immunofluorescence, also supported the specificity of our staining (Supplementary Figure 1e-g). Thus we conclude that the APC/C is localized to or around spindle poles in mitosis.

It is tempting to speculate that APC/C molecules localized at spindle poles could be part of the centrosome. Indeed, purification of centrosomes from cells arrested in mitosis indicated co-elution of the APC/C with centrosomal fractions (Figure 1b). Comparison of centrosomes purified from mitotic and interphase cells showed a significant increase in the APC/C signal in mitotic cells, indicating that the APC/C localized to centrosomes specifically in mitosis (Figure 1c, d). Additionally, we also detected the APC/C co-activators Cdc20 and Cdh1, found previously to localize to the spindle poles (Kallio et al., 2002; Meghini et al., 2016; Zhou et al., 2003). We also detected BubR1 in centrosomal fractions, but did not observe co-elution of the other SAC proteins Mad2 or Bub3 (Figure 1b). This is in agreement with a previous observation that BubR1 localizes to centrosomes independently of the mitotic checkpoint complex (MCC) (Izumi et al., 2009).

**Fig 1:**
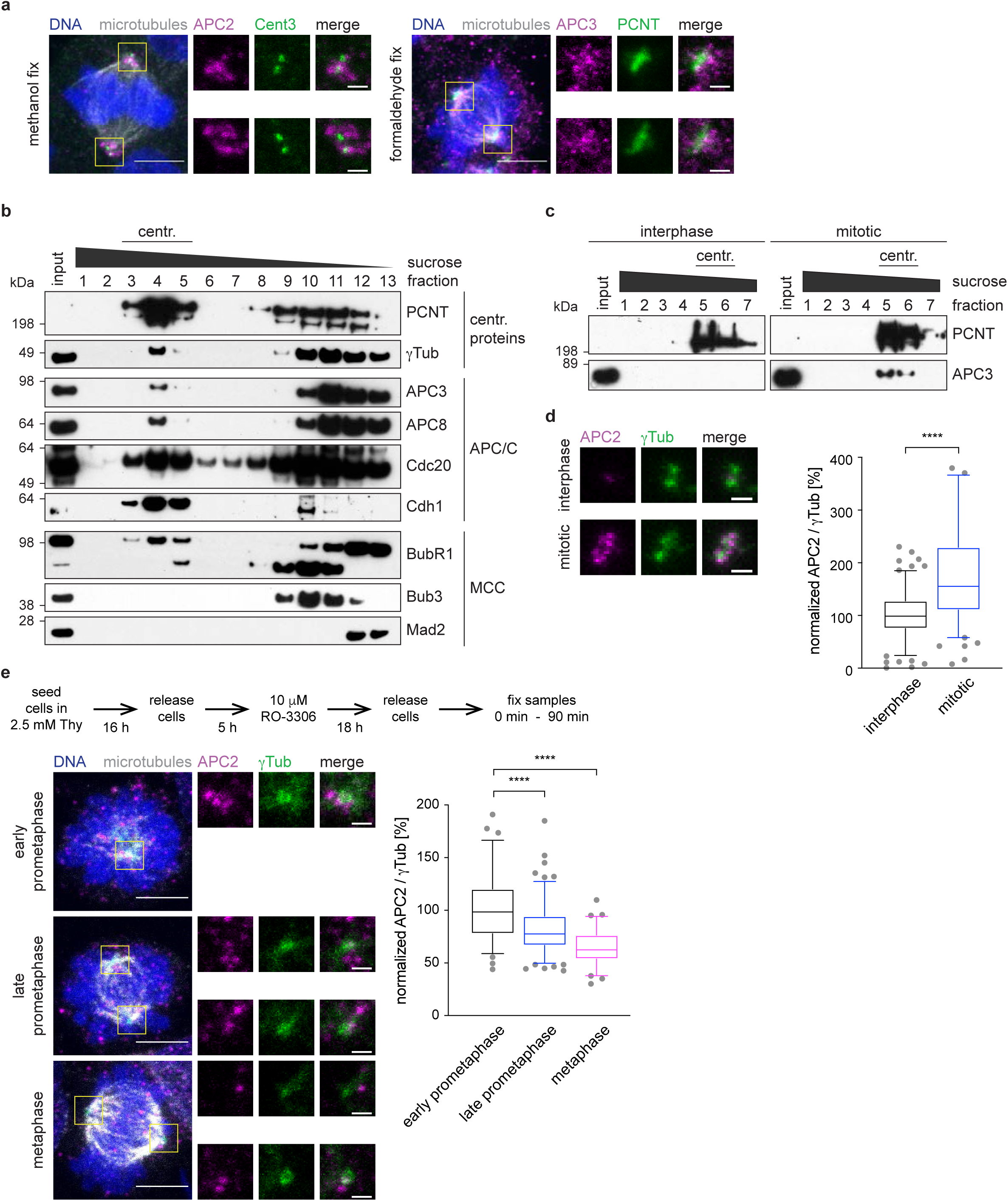
The APC/C is a component of the mitotic centrosome. **a**, Mitotic HEK293 cells were pre-extracted, fixed in ice cold methanol (left) or formaldehyde (right) and stained with the indicated antibodies. Scale bars are 5 μm in the overview and 1 μm in the inset. **b**, Centrosomes were purified from mitotic cells using a sucrose gradient. The eluted fractions were immunoblotted against the indicated proteins. The top shows PCNT and γTub as centrosomal proteins, the middle shows several APC/C components and MCC proteins are shown at the bottom. **c**, Centrosomes were purified in parallel from interphase cells and mitotic cells using a sucrose gradient. The first elution fractions were blotted against PCNT as a centrosomal marker and APC3. **d**, Centrosomes from **c** were fixed on a coverglass and stained with the indicated antibodies. The intensity of the APC/C was measured and normalized against the signal intensity on interphase centrosomes (right). **e**, Cells were treated according to the time line (top) and stained against the indicated proteins. The fluorescence intensity of APC2 was measured and normalized against the intensity in prophase. Scale bars are 5 μm in the overview and 1 μm in the inset. N = 77 (prophase), 130 (prometaphase), 73 (metaphase). ****P < 0.0001 using a simple one-way ANOVA test with Dunnett’s multiple comparison.

We observed that the APC/C is highly accumulated at the centrosome during early prometaphase, when cells possess mostly a monopolar spindle. With further progression through mitosis (late prometaphase) the APC/C signal at the centrosome decreased, being nearly undetectable in metaphase (Figure 1e).

### The APC/C localizes inside the PCM towards the proximal end of the centriole

To better understand the localization of the APC/C at the centrosome, we performed direct Stochastic Optical Reconstruction Microscopy (dSTORM) (Betzig et al., 2006; Rust et al., 2006) of prophase-arrested cells. In two dimensions, the APC/C localized in a regular, spot-like, circular pattern with a diameter of 524 +/-77 nm that is reminiscent of the 9-fold symmetry of centrioles (Supplementary Figure 2a, b). The interphase PCM consists of different layers or rings (Sonnen et al., 2012; Watanabe et al., 2019; Woodruff et al., 2014) and based on previous measurements, this placed the APC/C towards the outer region of the intermediate PCM, close to proteins such as Cep152, Cdk5rap2, and PCNT (Lawo et al., 2012; Sonnen et al., 2012). Using two-colour 3D-STORM, the APC/C was observed to accumulate mostly in one plane along the centriolar lumen marker centrin (Fong et al., 2014; Paoletti et al., 1996) (Supplementary Figure 2c and Supplementary Movie S1, S2). This indicated that the APC/C strongly localized either at the proximal or distal end of centrioles. Two-colour 3D STORM using the proximal end protein C-NAP1 showed that the APC/C and C-NAP1 reside within a similar Z-plane at the proximal end (Supplementary Figure 2d and Supplementary Movie S3, S4). Analysis of single centrosomal APC/C dots showed a diameter of 76 +/-28 nm (Supplementary Figure 2f), which corresponds closely to the calculated size of a single APC/C molecule of 35 nm plus two times 15 nm for the primary and secondary antibodies. To test this result, we used human APC/C purified from insect cells and performed dSTORM imaging using the same conditions. The measured diameter in this case was 69 +/-20 nm (Supplementary Figure 2e, f), indicating that the APC/C dots at the centrosome very likely represent only one or a maximum of two APC/C molecules. In conclusion, we show that single APC/C molecules accumulate in a symmetrical ring-like pattern surrounding the proximal end of the centriole within the intermediate PCM (Supplementary Figure 2g).

### The APC/C interacts with several centrosomal proteins

To understand which protein(s) are responsible for targeting the APC/C to the centrosome and to determine if the APC/C is directly involved in mitotic spindle assembly, we performed proximity labelling and mass-spectrometry. For this, we tagged the APC/C subunits APC2 and APC3 in independent cell lines with BioID2 under the control of a tetracycline inducible promotor (Supplementary Figure 3a). This improved version (Kim et al., 2016) of the biotin ligase BirA is constitutively active. Immunoprecipitation from whole cell lysates using streptavidin confirmed the successful incorporation of the BioID2-tagged APC/C subunits into the APC/C, as well as the proficient labelling with biotin (Supplementary Figure 3b). To specifically enrich for centrosomal proteins that interact with the APC/C, we purified centrosomes from our BioID2 cell lines (Supplementary Figure 3c). We noticed that most of the endogenous APC2 in our centrosomal fractions is replaced by the BioID-tagged protein, suggesting efficient incorporation and subsequent labelling with biotin (Supplementary Figure 3d). The fraction of BioID-tagged APC3 found at the centrosome seems to be lower compared to APC2 and the subsequent streptavidin pulldown confirmed weaker biotin labelling of these samples (Supplementary Figure 3c, d). The streptavidin pulldown from the purified centrosomes was then analysed by mass spectrometry. Label-free quantification (Zybailov et al., 2006) of the different datasets identified several centrosomal proteins that showed a strong enrichment in both tagged cell lines (Supplementary Figure 3e and Supplementary Table 1). These included Cep131, Cep152, Cep170, Cep192, Cep350 and PCM1. Subsequent analysis focused on the Cep proteins, but not PCM1, since it is the main component of pericentriolar satellites where the APC/C localizes during interphase (Supplementary Figure 1a). However, satellites are dissolved during mitosis, and PCM1 is not localized at the centrosome at this time (Kubo and Tsukita, 2003). It was therefore excluded as being a false positive, probably caused by potential contamination with interphase cells in our sample preparation.

To investigate the relationship between the identified proteins and the APC/C, we separately depleted the different Cep proteins (Figure 2a) and measured the intensity of the APC/C at the centrosome in mitosis (Figure 2b, 2c). Removal of most Cep proteins resulted in various degrees of reduced APC/C intensity, except for Cep350 where APC/C intensity at the centrosome remained unchanged (Figure 2b, 2c). Subsequent analysis focused on Cep131, Cep152, Cep170 and Cep192, excluding Cep350. Depletion of centrosomal proteins can result in reduction of microtubule nucleation (Gomez-Ferreria et al., 2007; Yan et al., 2006). This, however, is not the reason the APC/C does not localize to the centrosome anymore. Treatment of mitotic cells with the microtubule poison nocodazole showed that even in the absence of microtubules the APC/C still localized to the centrosome (Supplementary Figure 4a).

**Fig 2:**
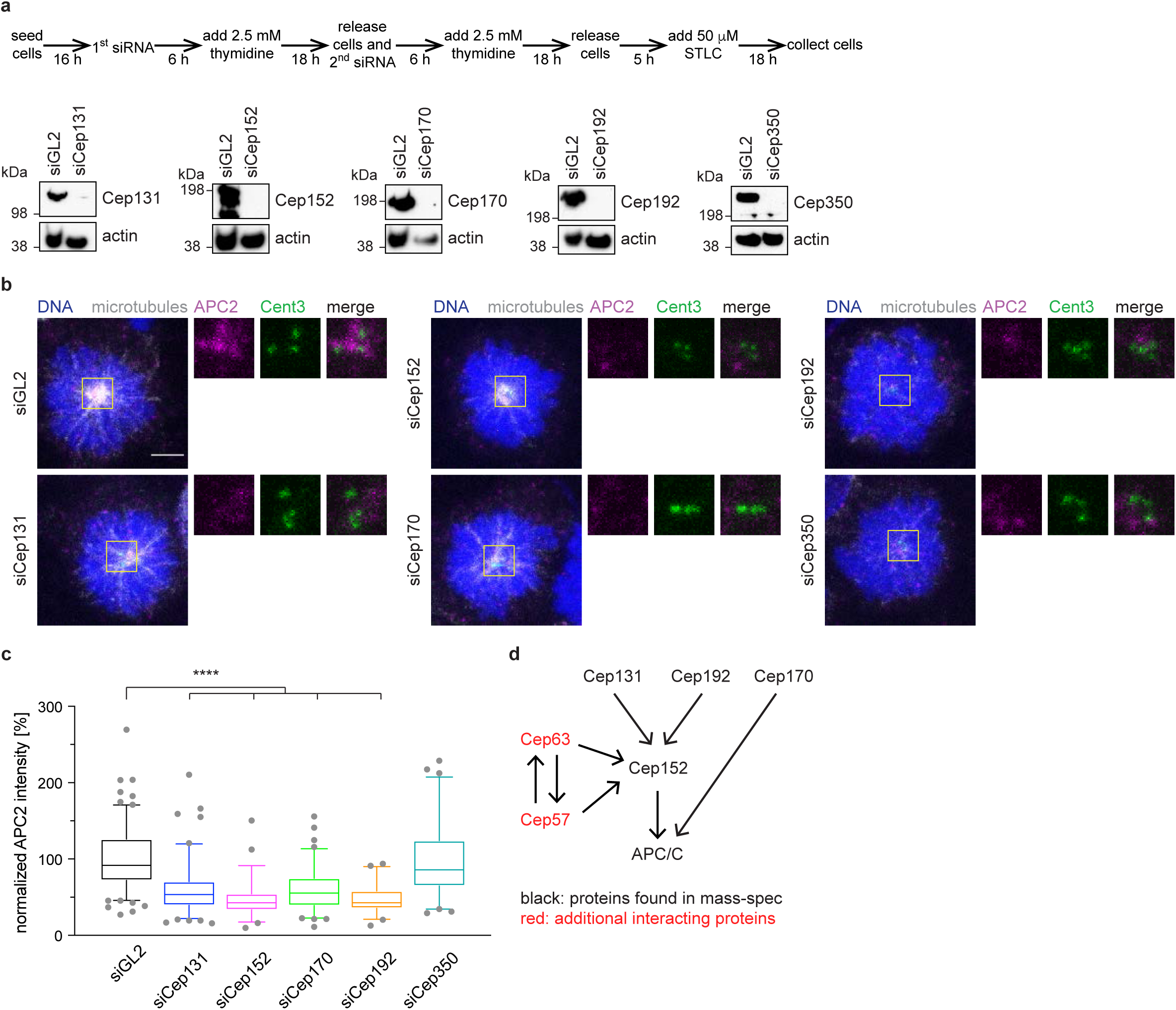
Depletion of centrosomal proteins leads to loss of the APC/C from the centrosome. **a**, HEK293 cells were depleted of the centrosomal proteins chosen for further analysis using siRNA according to the time schedule shown (top). STLC arrested, mitotic whole cell lysates were immunoblotted against the indicated proteins. Actin serves as a loading control. **b**, Cells were treated as in **a** and stained against the indicated proteins. **c**, Quantification of the APC/C intensity at the centrosome from cells shown in **b**. The intensity was normalized against the siGL2 control. N = 140 (siGL2), 109 (siCep131), 52 (siCep152), 86 (siCep70), 51 (siCep192), 63 (siCep350). ****P < 0.0001 using a simple one-way ANOVA test with Dunnett’s multiple comparison. **d**, Interaction model of the different centrosomal proteins with each other and with the APC/C.

Some of the Cep proteins are known to either interact with each other or depend on one of the other proteins for their localization (Kim et al., 2013; Kodani et al., 2015; Sonnen et al., 2013). To generate a small-scale interaction network, we measured the intensity of each Cep protein under the depletion conditions of the other proteins (Supplementary Figure 4b - e). As expected, the fluorescent intensity of each protein is reduced when it was depleted itself, confirming the specificity of the staining. Cep131, Cep170 and Cep192 localize independently to the centrosome and did not depend on any of the other proteins. However, Cep152 intensity at the centrosome was reduced in the absence of Cep131 and Cep192 (Supplementary Figure 4c), consistent with previous data (Kim et al., 2013; Kodani et al., 2015; Sonnen et al., 2013). It therefore seems likely that the reduced APC/C intensity upon depletion of Cep131 and Cep192 is a secondary effect due to the absence of Cep152 in these conditions (Figure 2e). Based on these data, indicating that Cep170 appeared to belong to a different interaction network, we excluded Cep170 from further analysis. Immunoprecipitation from cell lines expressing inducible eGFP-tagged Cep152 showed that it interacts with the APC/C (Figure 3f), validating our BioID and mass-spectrometry data. However, we cannot exclude that these interactions could also be indirect and mediated by the other centrosomal proteins. Additional Cep proteins, not detected in our mass-spectrometric analysis, can recruit Cep152 to the centrosome (Figure 2d). These include Cep57 and Cep63 that form a stable complex with Cep152 (Lukinavicius et al., 2013), as we will discuss later. In conclusion, Cep152 is the main, direct or indirect, interacting protein of the APC/C at the centrosome, with smaller contributions from other proteins.

**Fig 3:**
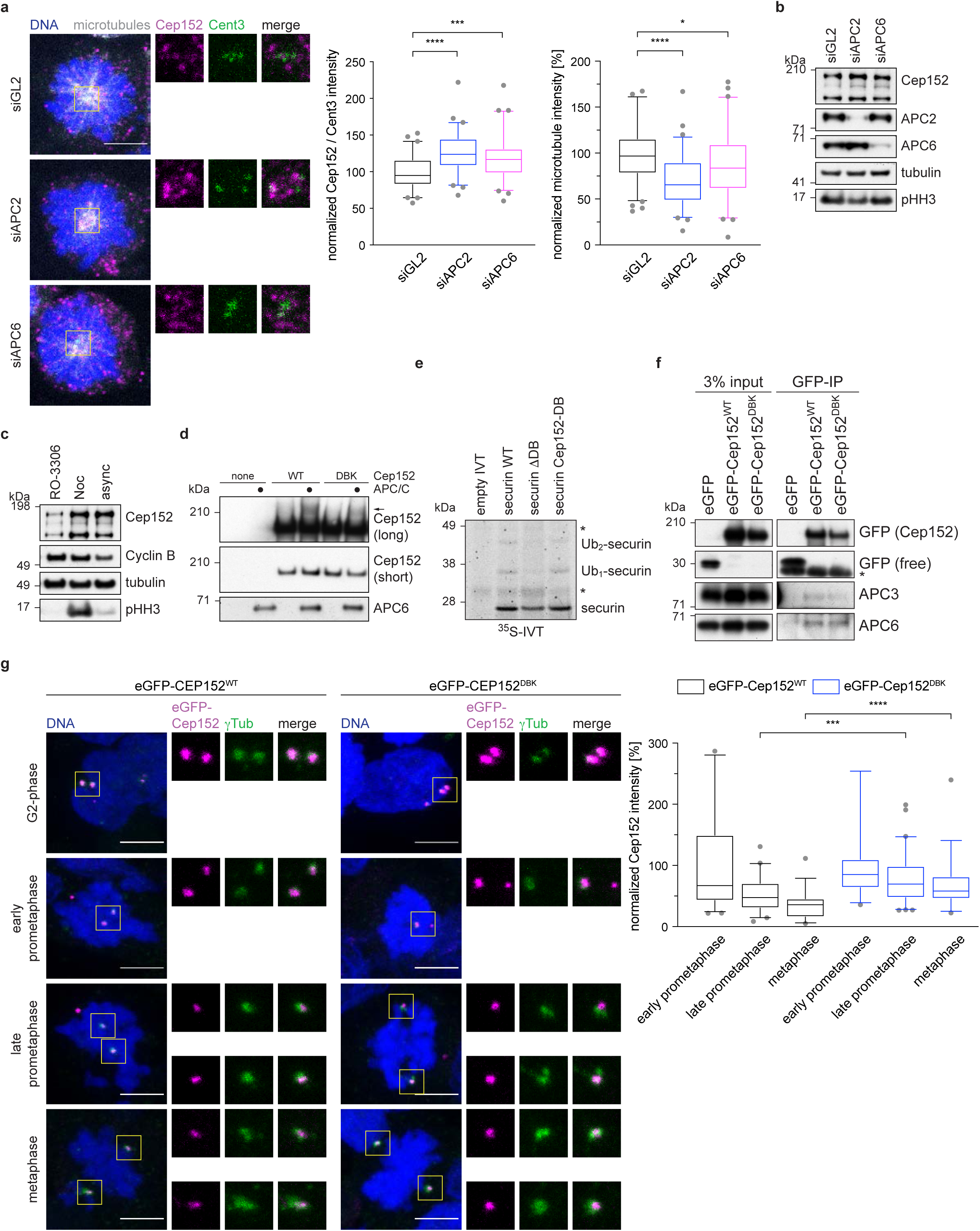
Cep152 is a substrate of the APC/C. **a**, HEK293 cell were treated with siRNA against APC2, APC6 or GL2 (control) and arrested in mitosis with STLC. Cells were stained against the indicated proteins. The fluorescence intensity of Cep152 at the centrosome was measured and normalized against the GL2 control (left). The fluorescence intensity of microtubules was measured and normalized against the GL2 control (right). N = 60 (siGL2), 60 (siAPC2), 62 (siAPC6). *P = 0.05, ***P = 0.0002, ****P < 0.0001 using a simple one-way ANOVA test with Dunnett’s multiple comparison. **b**, HEK cells were treated as described in **a** and whole cell lysates were immunoblotted against the indicated proteins. **c**, HEK cells were treated with the indicated compounds (Noc = nocodazole, async = asynchronous cells) and whole cell lysate was probed against the indicated proteins. **d**, eGFP-Cep152 was immunoprecipitated from HEK cells expressing the indicated constructs. The eluate was used for in vitro ubiquitination assays by incubation with purified APC/C. Arrow marks the ubiquitinated band. **e**, Radio-labelled securin was incubated in vitro with purified APC/C. WT = wildtype, ΔDB = deletion of the securin D-box, Cep152-DB = the sequence surrounding the securin D-box was replaced with the Cep152 D-box sequence (ATRKALGTVNR → QLRSELDKLNK). Asterisk marks unspecific bands. **f**, EGFP-tagged Cep152 proteins were immunoprecipitated from HEK cells expressing the indicated proteins. The co-eluted proteins were probed by immunoblot. Asterisk marks unspecific bands. **g**, HEK293 FlpIn T-Rex cell lines expressing eGFP-Cep152 wildtype (WT) or D box/KEN box (DBK) mutant proteins were treated as shown in **Fig 1e** and stained against the indicated proteins (left). The fluorescence intensity of eGFP-Cep152 at the centrosome was measured and normalized against the intensity of prophase cells of the corresponding cell line (right). N = 44 (WT, prophase), 34 (DBK, prophase), 41 (WT, prometaphase), 61 (DBK, prometaphase), 33 (WT, metaphase), 38 (DBK, metaphase). ***P = 0.0005, ****P < 0.0001 using a Mann-Whitney U-test.

### Cep152 is a substrate of the APC/C

Curiously, upon depletion of the APC/C by siRNA, Cep152 protein levels detected by immunofluorescence at the centrosome increased, while at the same time the intensity of spindle microtubules decreased (Figure 3a). This was accompanied by a small increase in total Cep152 levels as shown by immunoblotting (Figure 3b). Cep152 protein levels were also increased in the presence of the microtubule poison nocodazole that activates the spindle assembly checkpoint and therefore inhibits the APC/C (Figure 3c and Supplementary Figure 4a). These observations prompted us to investigate whether Cep152 is not only responsible for targeting the APC/C to the centrosome, but is also a substrate of the APC/C. Staining of cells during different stages of mitosis (early prometaphase to metaphase) showed that the intensity of Cep152 at the centrosome decreased (Supplementary Figure 5a) in a similar fashion as the APC/C intensity (Figure 1e). Sequence analysis (Jehl et al., 2016) of Cep152 revealed a potential D box and KEN box pair, the degron recognition motifs of APC/C substrates, present between residues 716 and 742 (Supplementary Figure 5b). We mutated the putative D box and KEN box together (called DBK mutant) and created cell lines that inducibly express eGFP-tagged variants of Cep152 (Supplementary Figure 5d). These eGFP-tagged Cep152 variants supported the established function of Cep152 during centriole duplication (Blachon et al., 2008) as they are able to rescue the absence of the endogenous protein (Supplementary Figure 5f). Due to the instability of recombinant Cep152, we were unable to both assess direct binding to the APC/C and assay APC/C-mediated ubiquitination of recombinant Cep152 *in vitro*. However, to test whether Cep152 is a potential APC/C substrate, we immunoprecipitated eGFP-tagged Cep152 from our HEK cell lines and used these proteins in an *in vitro* ubiquitination assay with purified APC/C. A portion of eGFP-Cep152^WT^ (wild-type) was shifted to a higher molecular weight species on incubation with the APC/C. This Cep152 modification depended on the APC/C and the D- and KEN boxes, being mostly ablated in the eGFP-Cep152^DBK^ mutant (Figure 3c), consistent with APC/C-catalysed Cep152 ubiquitination. To further investigate if Cep152 is an APC/C substrate, the D box of securin was replaced with the sequence of the identified Cep152 D box. Using IVT-generated ^35^S-labelled-securin in an *in vitro* ubiquitination assay showed that both wild type securin, and the mutant securin substituted with the putative Cep152 D box, were ubiquitinated to similar extents, whereas removing the D box of securin almost completely abolished ubiquitination (Figure 3d). This indicated that Cep152 contains a transferable D box, a hallmark of bona fide APC/C substrates (Zhang et al., 2019). Additionally, a small peptide that comprises the sequence of Cep152 incorporating the D box/KEN box pair was tested in an ubiquitination-competition assay. Ubiquitination of cyclin B1 was abolished in the presence of a peptide modelled on the well-established APC/C substrate Hsl1, and drastically reduced when the Cep152 peptide was used (Supplementary Figure 5e). These data support the proposal that Cep152 is an APC/C substrate, thereby explaining why levels of Cep152 increase when the APC/C is either depleted or inactivated (Figure 3a, b). Interestingly, as shown by co-immunoprecipitation, mutation of the Cep152 D box and KEN box does not abolish the binding of Cep152 to the APC/C (Figure 3f).

We then analysed the localization of eGFP-Cep152 during different mitotic stages. EGFP-Cep152^WT^ served as a control. Similar to the endogenous protein, its levels at centrosomes reduced as cells progressed from early prometaphase to metaphase (Figure 3g and Supplementary Figure 5a). As expected, both eGFP-tagged proteins localized to the centrosome in G2-phase (Figure 3g). Notably, compared with eGFP-Cep152^WT^, the centrosomal levels of eGFP-Cep152^DBK^ decreased less markedly as cells progressed through mitosis (Figure 3g). When cells were released from a G2 arrest into mitosis, unlike cyclin B1, Cep152 levels did not decrease (Supplementary Figure 5c). Taken together with our results that both APC/C depletion, and activation of the SAC, only slightly stabilizes Cep152 (Figure 3b, c), these data suggest that the APC/C regulates Cep152 levels either locally at the centrosome or by displacing Cep152 from the centrosome.

### Cep152 is an inhibitor of mitotic spindle assembly

Having established that Cep152 is responsible for recruiting the APC/C to the centrosome, and is itself a substrate of the APC/C, we next asked whether the APC/C-regulated microtubule nucleation (Figure 3a, b) is mediated through Cep152. A mitotic role for Cep152 has not previously been established. Cells expressing eGFP-Cep152^DBK^ exhibited a strong reduction in spindle microtubule intensity compared to cells expressing eGFP-Cep152^WT^ (Figure 4a), reminiscent of cells where the APC/C was depleted (Figure 3a, b). To further investigate this observation, a cold treatment to depolymerize all microtubules was performed in mitotic cells. After shifting cells back to 37 °C, nucleation of new microtubules from the centrosome was measured at different time points. In cells expressing eGFP-Cep152^WT^, microtubules reappeared after 5 minutes. In contrast, cells expressing eGFP-Cep152^DBK^ showed both a delay in microtubule nucleation (Figure 4c), and in establishing a bipolar spindle (Supplementary Figure 6a). This confirmed that the proper removal of Cep152 from the centrosome by the APC/C is a critical step for microtubule nucleation.

**Fig 4:**
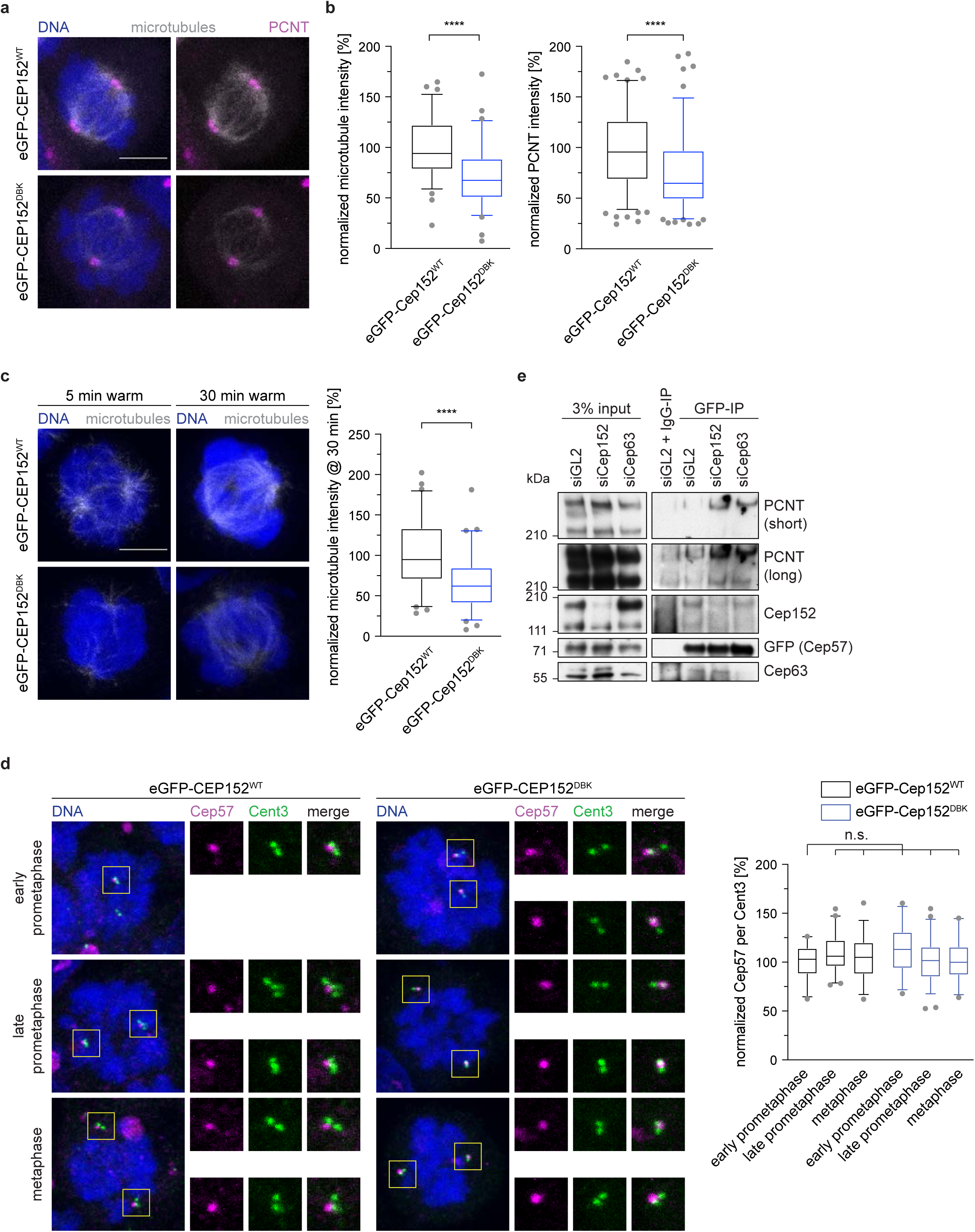
Cep152 is an inhibitor of mitotic spindle assembly and forms a complex with Cep57. **a**, HEK293 FlpIn T-Rex cell lines expressing eGFP-Cep152 wildtype (WT) or D box/KEN box (DBK) mutant proteins were treated as shown in **Fig 1e** and stained against the indicated proteins. **b**, The fluorescence intensity of tubulin in cells from **a** was measured in the whole cell and normalized against the eGFP-Cep152^WT^ cell line (left). N = 78 (WT), 77 (DBK). ****P < 0.0001 using a simple one-way ANOVA test with Dunnett’s multiple comparison. The fluorescence intensity of PCNT at the centrosome in cells from **a** was measured and normalized against the eGFP-Cep152^WT^ cell line (right). N = 158 (WT), 154 (DBK). ****P < 0.0001 using a simple one-way ANOVA test with Dunnett’s multiple comparison. **c**, HEK293 FlpIn T-Rex cell lines expressing eGFP-Cep152 wildtype (WT) or D box/KEN box (DBK) mutant proteins were subjected to 45 min at 4 °C. Cells were fixed at the indicated times after shifting them back to 37 °C and stained against tubulin (left). The fluorescent intensity of tubulin was measured in the whole cell at the 30 min after shifting cells back to 37 °C and normalized against the eGFP-Cep152^WT^ cell line (right). N = 61 (WT), 66 (DBK). ****P < 0.0001 using a simple one-way ANOVA test with Dunnett’s multiple comparison. **d**, HEK293 FlpIn T-Rex cell lines expressing eGFP-Cep152 WT or DBK mutant protein were treated as shown in **Fig 1e** and stained with the indicated proteins (left). The fluorescence intensity of Cep57 was measured around the centrosome and normalized against prophase of the eGFP-Cep152^WT^ cell line (right). N = 23 (prophase, WT), 48 (prometaphase, WT), 33 (metaphase, WT), 26 (prophase, DBK), 59 (prometaphase, DBK), 26 (metaphase, DBK). n.s. = not significant using a simple one-way ANOVA test with Dunnett’s multiple comparison. **e**, HEK cells expressing eGFP-tagged Cep57 were treated with control siRNA (siGL2), siRNA against Cep152 (siCep152), or siRNA against Cep63 (siCep63) and lysed for anti-GFP immunoprecipitation. The eluates were probed against for the indicated proteins by immunoblotting.

Since Cep152 is not known to directly interact with microtubules, we speculated that additional interactions partners of Cep152 might be involved in this process. Cep152 forms a stable complex with two other centrosomal proteins, Cep57 and Cep63 (Lukinavicius et al., 2013; Watanabe et al., 2019). In this trimeric complex, Cep63 bridges the interaction between Cep57 and Cep152 (Wei et al., 2020; Zhao et al., 2020). Nucleation of microtubules in mitosis depends on the successful accumulation and expansion of the PCM, comprising proteins such as PCNT and Cdk5rap2 (Gould and Borisy, 1977; Woodruff et al., 2017). Since Cep57 recruits PCNT to centrosomes (Watanabe et al., 2019), we tested if Cep152 controls centrosomal PCNT levels. Notably, we found that the levels of PCNT at centrosomes were reduced in the presence of stabilized eGFP-Cep152^DBK^ (Figure 4a, b). We therefore speculated that during mitosis, release of Cep57 from the Cep152-Cep63-Cep57 complex allows it to bind PCNT. We found that Cep57 levels at the centrosome remain constant throughout mitosis (Figure 4d) in contrast to Cep152 and Cep63 that both decreased (Supplementary Figure 5a, Supplementary Figure 6b). In agreement with our proposal, in the presence of stabilized eGFP-Cep152^DBK^, Cep63 remains at the centrosome (Supplementary Figure 6b). These findings suggested the possibility that once Cep152 and Cep63 are removed from the centrosome, Cep57 is free to recruit additional PCNT to enhance microtubule nucleation. To test our hypothesis directly, we separately depleted Cep152 and Cep63 and performed an immunoprecipitation of eGFP-tagged Cep57 from mitotic cells. The immunoprecipitation confirmed previous data that Cep63 bridges the interaction of Cep152 with Cep57 (Watanabe et al., 2019; Wei et al., 2020; Zhao et al., 2020), since in the absence of Cep63 less Cep152 was bound to eGFP-Cep57. In contrast, depletion of Cep152 did not reduce the binding of Cep63 to eGFP-Cep57 (Figure 4e). Cep63 is difficult to deplete in HEK cells. About 50% of the protein remained even after 96 hours of siRNA treatment. Nevertheless, upon siRNA mediated reduction of either Cep152 or Cep63 more PCNT is associated with eGFP-Cep57 (Figure 4e), consistent with the hypothesis that the Cep152-Cep63 complex inhibits Cep57 – PCNT interactions. In conclusion, the removal of Cep152 from the centrosome by the APC/C during mitosis is important to liberate Cep57 from its inhibitory complex and aid faithful microtubule nucleation.

### Persistent Cep152 at the centrosome leads to mitotic errors

Based on our data that microtubule nucleation and formation of a bipolar spindle are slowed down in the presence of eGFP-Cep152^DBK^ (Figure 4c, Supplementary Figure 6a), we reasoned that cells expressing stabilized eGFP-Cep152^DBK^ would need more time to complete mitosis. We performed live cell microscopy imaging of cells expressing the different eGFP-Cep152 variants and analysed the time from nuclear envelope breakdown (NEBD) to metaphase, and the time from metaphase to anaphase onset. In cells expressing eGFP-Cep152^DBK^ both phases were prolonged compared to cells with eGFP-Cep152^WT^ (Figure 5a, b), consistent with our data that in these cells microtubule nucleation is diminished. An increased time from metaphase to anaphase onset indicates that these cells might have problems with correctly attaching their chromosomes to the mitotic spindle, resulting in mitotic errors. Consistent with this, we found that the number of cells that showed either mis-aligned chromosomes during metaphase, or lagging chromosomes during anaphase (together referred to as “mitotic errors”), was increased with eGFP-Cep152^DBK^ (Figure 5c and Supplementary Movies S5, S6). These data highlight the crucial role of APC/C-mediated Cep152 degradation during mitosis, which is required to perform successful spindle assembly.

**Fig 5:**
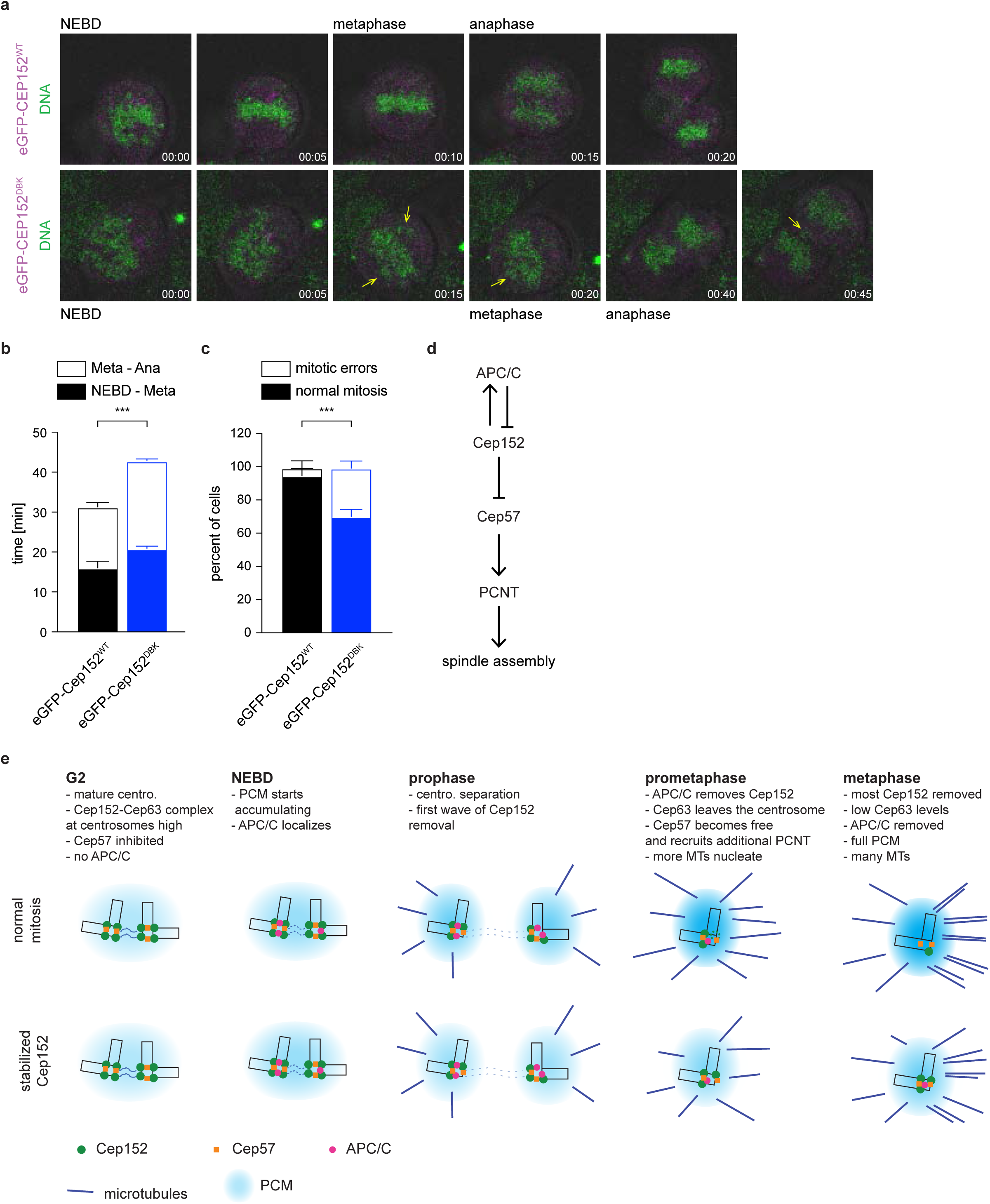
Persistent Cep152 at the centrosome leads to mitotic errors. **a**, eGFP-Cep152^WT^ and eGFP-Cep152^DBK^ cell lines were subjected to live cell imaging. Exemplary still images from the movies are shown. See Supplementary Movie 5 and 6. Nuclear envelope breakdown (NEBD), metaphase and anaphase onset are marked. DNA was stained with SiR-DNA. Unaligned chromosomes are indicated with yellow arrows. **b**, Cells from **a** were analysed for the time they need to progress from NEBD to metaphase (NEBD – Meta) and from metaphase to anaphase onset (Meta – Ana). Bar graphs show the mean +/-SEM. ***P = 0.0006 using a simple one-way ANOVA test with Dunnett’s multiple comparison. N = 4 experiments, WT = 197 cells, DBK = 189 cells. **c**, Cells from **a** were analysed for misaligned chromosomes during metaphase and lagging chromosomes during anaphase, combined as mitotic errors. Bar graphs show the mean +/-SEM. ***P = 0.0005 using a Holm-Sidak t-test. N = 4 experiments. **d**, Regulatory network of the APC/C and centrosomal proteins that regulate mitotic spindle assembly. The main APC/C-interacting protein at the centrosome is Cep152, which is also an APC/C substrate creating a negative feedback loop. **e**, Model of microtubule nucleation in the presence of wild type (top) or stabilized (bottom) Cep152.

## Discussion

This study reveals the crucial role of the APC/C at the centrosome. Recruitment of the APC/C to the centrosome depends on Cep152, which is itself an APC/C substrate. This negative feedback loop ensures that Cep152 auto-regulates its removal from the centrosome, a step that is important for proper microtubule nucleation mediated by Cep57 and PCNT (Figure 5d, 5e).

Based on our data we propose that during interphase the APC/C accumulates in centriolar satellites. Previous studies had shown that the APC/C is localized to spindle poles in human cells and that this localization depends on the proteins NuMA and dynein (Ban et al., 2007), both of which have been implicated in the transport of centriolar satellites (Engelender et al., 1997; Merdes et al., 2000; Merdes et al., 1996). Indeed, a recent proteomic study identified the APC/C as a minor component of centriolar satellites (Gheiratmand et al., 2019). This suggests the possibility that the APC/C is a dynamic component of centriolar satellites and is transported to the centrosome together with some of its interacting proteins. It also offers an explanation for the Cep170 depletion phenotype that leads to a reduction of APC/C intensity at the centrosome. Cep170 was recently identified as a component of centriolar satellites (Barenz et al., 2018; Gheiratmand et al., 2019; Gupta et al., 2015; Quarantotti et al., 2019). Its centrosomal localization was shown to be mainly at the distal end of the mother centriole (Guarguaglini et al., 2005; Mazo et al., 2016), where it cannot interact with the APC/C, which we show localizes proximally. However, if satellite-localized Cep170 is involved in transporting the APC/C to the centrosome, Cep170 depletion could result in a failure to do so. A similar effect could also contribute to the Cep131 depletion phenotype where the APC/C is also absent from centrosome. However, in this instance a direct relationship between Cep131 and Cep152 was previously established (Kodani et al., 2015) and confirmed by our study, which we propose to account for the main phenotype.

Once satellites disassemble in mitosis (Kubo and Tsukita, 2003) the APC/C localizes to the centrosome inside the PCM. This behaviour is similar to other proteins such as PCNT, Cdk5rap2 and γTub, which, as for the APC/C, reside in a similar layer surrounding the centrosome (Lawo et al., 2012; Sonnen et al., 2012). The main interacting protein of the APC/C at the centrosome is Cep152. Cep152 together with its interacting proteins Cep63 and Cep57 have previously been shown to localize to the proximal end of the centriole, forming a ring-like pattern (Lukinavicius et al., 2013; Sonnen et al., 2012; Zhao et al., 2020). This is in agreement with our data showing a similar localization for the APC/C. Our data establish that in the absence of Cep152, APC/C can no longer localize properly to the centrosome. This localization, however, is important, as interfering with the timely removal of Cep152 from centrosomes by the APC/C results in reduced microtubule nucleation and consequently in chromosome alignment defects and segregation errors. These phenotypes are very similar to Cep57 depletion (Watanabe et al., 2019; Wu et al., 2012) and we established that the Cep152-Cep63 complex has an inhibitory function on Cep57. In the presence of stabilized Cep152, Cep63 remains at the centrosome and Cep57 is not released from this inhibition, which hampers the recruitment of PCNT.

A function of Cep152 in mitotic spindle assembly has not been previously established, probably because a stabilized Cep152 mutant was not available. Our work, therefore, also implicates Cep152 as an inhibitor of mitotic spindle formation. We postulate that to ensure the timely removal of Cep152 from the centrosome it auto-regulates its local ubiquitination by the APC/C. This very elegant dual function of Cep152 as a recruiter as well as a substrate of the APC/C ensures that Cep152 is ubiquitinated at centrosomes in a local and timely fashion, and not throughout the cell. We show that Cep152 accumulates only slightly throughout the cell in the absence of APC/C activity. This suggests that ubiquitination of Cep152 by the APC/C could result in relocalization of Cep152 from the centrosome, but not necessarily in its degradation. This behaviour has not been reported previously for APC/C substrates but is known in other contexts (Huang et al., 2003; Kruse and Gu, 2009; Yao et al., 2018). Our data do not exclude that there are additional APC/C substrates and interacting proteins localized to the centrosome. For example, the kinase Nek2A predominantly localizes to the centrosome in early mitosis until it is degraded by the APC/C (Faragher and Fry, 2003; Fry et al., 1998). Recently, it was shown that the *Drosophila* homologue of Cep192 (Spd2) recruits the APC/C co-activator Cdh1 to the centrosome and that Cep192/Spd2 is also an APC/C substrate (Meghini et al., 2016; Raff et al., 2002). This observation is supported by our data that we find Cdh1 co-eluting with isolated centrosomes and we also detected Cep192 in our APC/C proximity labelling. It is therefore possible that different parts of the APC/C, such as the core-complex and the co-activators, are localized to the centrosome by separate processes and a functional complex is only assembled when needed. This could favour a rapid activation of the APC/C at the centrosome to ensure that Cep152 is targeted early in mitosis, because persistent Cep152 delays microtubule nucleation. The details of the mechanisms responsible for regulating the APC/C localized to centrosomes are not currently understood.

Taken together, our study establishes how the APC/C is recruited to the centrosome and reveals its critical function to locally ubiquitinate Cep152, an inhibitor of mitotic spindle assembly. It also shows that the APC/C has a dual function during mitosis, where it is not only involved in mitotic progression, but also directly regulates spindle assembly.

## Material and Methods

### Tissue culture and cell cycle synchronization

HEK293 FlpIn-TRex cells (Invitrogen) were cultured in DMEM (Gibco) supplemented with 10% tetracycline free FBS (PAN Biotech) at 37 °C and 5% CO_2_. For the stable integration of eGFP-Cep131, eGFP-Cep192, eGFP-Cep152 (WT, DBK), eGFP-APC2 and APC3-eGFP into the genome the corresponding genes were cloned into the pcDNA5-FRT-TO vector (Invitrogen) with an N-terminal or C-terminal (only in the case of APC3) eGFP tag. HEK293 FlpIn-TREX cells were co-transfected with the respective pCDNA5-FRT-TO plasmid and pOG44, containing the flp-recombinase, using HBS buffer. Briefly, cells were seeded the evening before transfection and the medium was exchanged the next morning. Both plasmids were mixed with 160 mM CaCl_2_ and 2x HBS buffer (final concentrations: 137 mM NaCl, 5 mM KCl, 0.7 mM Na_2_HPO_4_, 7.5 mM D-glucose, 21 mM HEPES) and added to the cells. The next steps were performed according to the Invitrogen manual. Cells were selected using 100 μg/mL Hygromycin B gold (InVivoGen). For creation of the BioID2-APC2 and APC3-BioID2 cell lines the same protocol was performed, but the pcDNA5-FRT-TO vector was modified to either include an N-terminal or a C-terminal BioID2 tag. All stable cell lines were kept under selection. Protein expression was induced with 200 pg/mL doxycyclin (Sigma). Cell cycle synchronization was performed using a combination of 2.5 mM thymidine (Sigma) to arrest cells in or before S-phase and either 300 nM nocodazole (Sigma) or 500 nM taxol (Sigma) or 5 μM STLC (Sigma) or 10 μM RO-3305 (Santa Cruz) for a mitotic arrest. Cells were released from thymidine after 16 h by washing three times with pre-warmed medium using the same volume as during culture. The second drug was added 4 h after thymidine release and incubated for another 16 h. Release from RO-3306 was performed in the same way.

### RNAi mediated protein depletion

Cells were seeded at a density of 50% 16 h before transfection with siRNA. RNAi was performed using RNAiMAX reagent (Invitrogen) according to the manual. For the depletion of all proteins a double depletion protocol using two times 20 nM oligo was used. In short, 16 h after seeding the cells they were transfected with the first round of siRNA and 8 h later thymidine was added (see above). Again 16 h later the cells were released from thymidine and the second round of siRNA transfection was performed. The thymidine block and release were repeated and cells were arrested in mitosis afterwards (see above). This protocol ensures that most of the cells only perform one round of mitosis, which avoids problems with centrosome duplication (in case of the depletion of centrosomal proteins) or cell cycle arrest (in case of APC/C depletion). The following siRNA oligos were used: GL2 AACGUACGCGGAAUACUUCGA[dt][dt], APC6_5 AUGAUGCUCUAGAUAACCGAA[dt][dt], APC6_6 CCCAUGCACUUCGGUCACGAA[dt][dt], APC2_6 CUCACUGGAUCGUAUCUACAA[dt][dt], APC2_5 AAGGUUCUUCUACCGCAUCUA[dt][dt], Cep131 CUGACAACUUGGAGAAAUU[dt][dt], Cep170 GAAGGAAUCCUCCAAGUCA[dt][dt], Cep350 AUGAACGAUAUCAGUGCUAUA[dt][dt], Cep192 AAGGAAGACAUUUUCAUCUCU[dt][dt], Cep152 GCGGAUCCAACUGGAAAUCUA[dt][dt], Cep63 GGCUCUGGCUGAACAAUCA[dt][dt]

### Immunoblotting

For immunoblotting about 3 × 10^6^ cells / mL were washed once with PBS, spun down and resuspended in NuPAGE LDS Sample Buffer (Invitrogen). After heating the solution for 5 min at 95 °C, five to ten microliters were loaded on to a Novex Bis-Tris 4% - 12% gel (Invitrogen) and run for the appropriate time. The proteins were transferred to a nitrocellulose membrane (GE Healthcare), which was blocked with 5% dried milk in PBS-Tx (PBS, 0.1% Triton-× 100). The primary antibodies were incubated over night at 4 °C, the secondary antibodies were used for 1 h at room temperature. The following antibodies were used for immunoblotting: pericentrin (Abcam ab4448, 1:2000), APC8 (Abcam ab182003 1:1000), APC6 (Cell Signaling 9499, 1:1000), APC3 (Cell Signaling 12530, 1:1000), APC2 (Cell Signaling 12301, 1:1000), CDH1 (Abcam ab89535, 1:1000), BubR1 (Abcam ab54894, 1:500), Mad2(Abcam ab10691, 1:1000), APC3 (Sigma C7104, 1:500), gamma-tubulin (Sigma T6557, 1:1000), beta-actin (Santa Cruz sc-47778-HRP, 1:2000), alpha-Tubulin (BioRad MCA78G, 1:2000), Cdc20 (Santa Cruz sc-8358, 1:100), Bub3 (BD 811730, 1:500), Cep131 / AZI-1 (Bethyl A301-415A, 1:1000), Cep170 (Abcam ab72505, 1:500), Cep350 (Novus NB100-59811, 1:500), Cep152 (Bethyl A302-480A, 1:500), Cep192 (Bethyl A302-324A, 1:1000), Cyclin B1 (Abcam ab72, 1:1000), His (Takara/Clontech 631212, 1:1000), GFP (Roche 11814460001, 1:1000), Cep57 (Genetex GTX115931, 1:1000), Cep63 (Fanni Gergely lab, Cancer Research UK, Cambridge), secondary anti-rabbit (Thermofisher 31462, 1:10.000), secondary anti-mouse (Agilent P0260, 1:10.000), secondary anti-rat (Santa Cruz 2032, 1:10.000).

### Centrosome purification

Mitotic centrosomes were purified from 2 × 10^8^ HEK293 cells. Cells were synchronized with thymidine and nocodazole (see above). One h before harvesting the cells 1 μg/mL Cytochalasin D (sigma) was added. Mitotic cells were collected by shake off and gentle washing with medium. All subsequent steps were performed at 4 °C. Cells were washed once in PBS, once in 0.1x PBS + 8% sucrose and once in 8% sucrose in H_2_0. The cells were resuspended in 10 mL lysis buffer (1 mM Pipes, 0.1 mM EGTA, 0.1% 2-mercaptoethanol, 0.5% Triton-× 100, EDTA-free complete protease inhibitors (Roche)) and incubated for 10 min. The lysate was centrifuged at 2500 g for 10 min and the supernatant filtered through a 40 μm cell strainer. From a 50x stock of PE (500 mM Pipes, 50 mM EGTA, pH 7.2) the respective volume to prepare a 1x solution was added to the supernatant. Sucrose solutions with 70% (w/v), 60% (w/v), 50% (w/v), and 40% (w/v) were prepared in 1x PE. The cell lysate was carefully layered over 1 mL 60% sucrose cushion in a thin walled Beckman tube (344058) and centrifuged at 10,000 g for 30 min using a slow deceleration. About 70% of the tube content was removed from the top and discarded and the remaining volume was mixed with the 60% sucrose cushion to get solution of about 20% sucrose. In parallel, a sucrose gradient consisting of 2 mL 70% sucrose (bottom), 1.2 mL 50% sucrose (middle) and 1.2 mL 40% sucrose (top) was prepared in a Beckman tube (344060). The cell lysate containing 20% sucrose was layered over the sucrose gradient and centrifuged for 16 h at 100,000 g using a no break deceleration. The tube with the gradient was pierced at the bottom with an 18G needle and fractions of the following sizes were collected: F1-3 each 500 μL, F4-9 each 200 μL, F10-11 each 500 μL, F12-13 each 1 mL. Centrosome containing fractions were snap frozen in liquid nitrogen.

### Mass-Spectrometry

To prepare samples for mass-spectrometry, cells were treated as described above, but additionally incubated with 50 μM biotin (Sigma) for 24 h before harvesting. After centrosomes were purified, all fractions from the corresponding cell lines were combined and diluted with three times to volume of lysis buffer plus 0.1% SDS. The solution was sonified using a microtip with the following settings: 1 min time, 10 seconds on, 20 seconds off, 45% power. The sonified centrosomes were incubated with magnetic streptavidin beads (MyOne Streptavidin C1, ThermoFisher) over night. Washing was performed one time each in the following order: lysis buffer, SDS buffer (2% SDS in H_2_O), salt buffer (500 mM NaCl, 1% Triton-× 100, 1 mM EDTA, 50 mM HEPES, pH 7.6), Tris buffer (50 mM Tris, 50 mM NaCl, pH 7.6). Purified proteins were eluted from the streptavidin beads with Tris buffer plus 5 mM biotin. The whole samples were run on a NuPAGE SDS gel, stained with InstantBlue (Expedeon) and the each lane was cut into 10 equally sized pieces, which were send to mass-spectrometric analysis as described before (Zhang et al., 2016).

### Co-Immunoprecipitation

HEK cells were treated with siRNA or compounds for synchronization as described above. For eGFP-Cep152 immunoprecipitation, cells were lysed in the same way as described for the centrosome purification up until the addition of PE-buffer. For eGFP-Cep57 immunoprecipitation, cells were lysed in a Tris buffer (20 mM Tris pH 7.4, 50 mM NaCl, 5 mM EGTA, 2 mM MgCl_2_, 1 mM DTT, 0.5 % Triton-× 100, complete protease inhibitors) for 30 minutes on ice. In both cases, the lysate was centrifuged for 10 min at 2.500 g at 4 °C and the supernatant was carefully taken off. Subsequently, it was incubated with 5 μg GFP antibody (Roche) or normal mouse IgG (santa cruz) coupled to magnetic protein G beads (Invitrogen) for 3 - 4 h at 4 °C under constant agitation. The beads were washed three times with the corresponding lysis buffer (+ 150 mM NaCl in case of eGFP-Cep152) and eluted by cooking in NuPAGE LDS Sample Buffer (Invitrogen) with 5 mM DTT.

### Live cell microscopy

A SP8 confocal microscope (Leica) equipped with a heated environmental chamber, an argon laser, a 630 nm laser line and an APO CS2 40x/1.1 water immersion lens was used for live cell imaging. Live cell microscopy was performed as described before (Zhang et al., 2019). The imaged cells were analysed for the time of NEBD, the time when a metaphase plate was observed for the first time and the time of anaphase onset. Mitotic errors that were visible during the movies were noted down and analysed as well.

### Immunofluorescence microscopy and antibodies

After cells were grown in tissue culture plates, mitotic cells were collected by a gentle wash with medium and spun down onto polylysine (Sigma) treated cover glasses for 5 min at 650 g. Cells were fixed using either methanol or formaldehyde. Except for where noted cells were pre-extracted for 2 min at 37 °C using 0.2% Triton-× 100 in PHEM buffer (60 mM Pipes, 25 mM Hepes, 10 mM EGTA, 2 mM MgCl_2_, pH 7.5). For methanol, cells were incubated for 5 min at –20 °C with pre-cooled methanol. Afterwards cells were rehydrated with PBS and washed with PBS-Tx, before blocking with ABDIL (1% BSA, 0.1% Triton-× 100, PBS). For formaldehyde, cells were incubated two times for 5 min at 37 °C with pre-warmed 3.7% formaldehyde in PHEM buffer. Subsequently cells were washed in PBS-Tx and blocked in ABDIL. Primary antibody incubation was performed for 1 h at room temperature. After washing three times with ABDIL, the secondary antibodies were incubated for 1 h at room temperature, and washed again three times in ABDIL. If microtubules were stained, the process was repeated with an alpha-tubulin antibody (BioRad MCA78G, 1:2000) to minimize cross-reaction. In the last washing step 1 μg/mL DAPI (Sigma) was included. Primary antibodies used for staining were: pericentrin (Abcam ab4448, 1:2000), APC2 (Cell Signaling 12301, 1:1000), APC3 (sigma C7104, 1:500), gamma-Tubulin (sigma T6557, 1:1000), Cep131 / AZI-1 (Bethyl A301-415A, 1:1000), Cep170 (Abcam ab72505, 1:500), Cep350 (Novus NB100-59811, 1:500), Cep152 (Bethyl A302-480A, 1:500), Cep192 (Bethyl A302-324A, 1:1000), GFP (Abcam ab6556, 1:1000), Centrin3 (Novus H00001070-M01, 1:1000), Cep57 (Genetex GTX115931, 1:500), Cep63 (Millipore 06-1292,1:100). Secondary antibodies were pre-absorbed and Alexafluor coupled from Thermofisher. Alexa-488, Alexa-568, and Alexa-647 were used in all combinations against mouse, rabbit, or rat primary antibodies.

Images were acquired with an SP8 confocal microscope (Leica) quipped either with 405 nm, 488 nm, 568 nm and 633 nm laser lines or equipped with a White Light Laser and a 592nm STED laser. Both microscopes were used with an APO CS2 63x/1.4 oil immersion objective. Image resolution was set to 1024×1024 with a 3x zoom-in, 600 Hz scan speed and 2x line average. A Z-spacing of 0.23 μm was always used. Laser powers and detector gain were set for each primary antibody individually, but were kept constant between different experiments with the same antibody to ensure reproducibility.

### Image quantification and statistical analysis

Fluorescent images were analysed with ImageJ. Maximum intensity projections were created and a circular region of interest was used for the measurements. This could either be a small circle in case of centrosomes or a circle including the whole cell in case of tubulin measurements; the background was measured outside the cells. The integrated density was corrected for the area and the background. All values for the control condition in each experiment were averaged and this mean value was used to normalize all other conditions against the control using Excel (Microsoft). The normalized values were exported to Prism 8 (Graphpad) and plotted. All boxplots display the 5% to 95% range (whiskers), the 25% and 75% range (box) and the median (line). Bar graphs display the mean +/-standard deviation. Statistical analysis was performed in Prism 8 and the tests that were used are described in the corresponding figures legends. All experiments were repeated at least twice using biological replicates.

### 2D and 3D dSTORM

Direct-STORM was performed with a Nikon N-STORM equipped with 405 nm, 488 nm, 561 nm and 647 nm laser lines (Agilent/Keysight MLC 4008) using an APO TIRF 100x/1.49 oil immersion objective. Cells were cultured in an 8-well glass bottom slide (Ibidi) and fixed and stained as described above. The secondary antibodies used were coupled to Alexafluor-568 or Alexaflour-647 and were additionally fixed with 4% formaldehyde after the staining was concluded (post fix). Before imaging, the wells were incubated for 5 min with Tetraspeck Beads (Thermofisher, 1:100) and subsequently washed with PBS. The cells were imaged in switching buffer (100 U glucose oxidase (Sigma), 10.000 U catalase (Sigma), 8% glucose, 100 mM MEA (Sigma), pH 7.6) by filling the well of the ibidi slide to the top and covering it with a cover glass avoiding air bubbles. For 3D imaging a cylindrical lens was inserted into the light path. STORM was performed by inducing and observing blinking events of the Alexafluor secondary antibodies, usually 50.000 frames with 10 ms exposure were recorded. Pre- and post-STORM widefield images were acquired as well. Point fitting in 2D and 3D space was performed within the Nikon software. For 2D STORM the resulting images were directly saved from the Nikon software. For 3D STORM the fitted point coordinates were exported, transformed in ImageJ using ChriSTORM and imported using the ThunderSTORM ImageJ plugin. If necessary, two-colour images were manually aligned using the Tetraspeck beads as reference points with the “Align RGB planes” plugin in ImageJ. A gaussian normalization algorithm was used for visualization and the data were rendered in 3D using the ImageJ 3D viewer. The distance between two APC/C dots to estimate centrosomal diameter was measured using the “Peak finder” plugin in ImageJ. The size of APC/C dots around the centrosome and of purified APC/C was measured using a Full Width Half Max (FWHM) macro by John Lim in ImageJ.

### Ubiquitination assays

For radiolabelling of securin with ^35^S-Met, the T7 TnT Quick coupled in vitro transcription/translation system (Promega) was used according to the suppliers instructions.

EGFP-tagged Cep152 for the ubiquitination assay was immunprecipitated from nocodazole arrested HEK cells. For this, cells were harvested, washed in PBS and lysed in buffer (150 mM NaCl, 1 mM MgCl_2_, 50 mM HEPES pH 8, 1 mM EGTA, 0.5% Triton-× 100, complete protease inhibitors (Roche)). The lysate was centrifuged for 10 min at 2.500 g at 4 °C and the supernatant was carefully taken off. It was subsequently incubated with 5 μg GFP antibody (Roche) or normal mouse IgG (santa cruz) coupled to magnetic protein G beads (Invitrogen) for 2 h at 4 °C under constant agitation. The beads were washed three times with lysis buffer + additional 150 mM NaCl (for a total 300 mM NaCl) and eluted by incubation with glycine buffer (150 mM NaCl, 200 mM glycine, pH 2.3) for 10 min on ice. The solutionwas neutralized by addition of a corresponding volume of 4 M NaOH to reach pH 7.5.

In vitro ubiquitination was performed using APC/C and Cdc20 purified from insect cells. 60 nM APC/C, 30 nM Cdc20, 90 nM UBA1, 300 nM UbcH10 or 300 nM UbcH5, 35 μM ubiquitin, 5 mM ATP, 10 mM MgCl_2_, were mixed in a buffer containing 40 mM HEPES (pH 8.0), 80 mM NaCl, 0.6 mM DTT. The reaction was mixed with 10% radiolabeled IVT or 50% eluted Cep152 and incubated for 60 min at 23 °C. The assay was stopped by the addition of one volume of 2x concentrated NuPAGE LDS loading buffer.

## Acknowledgements

We thank F. Gergely from the Cancer Research UK Cambridge Institute for antibodies and discussions. M. van Breugel from the MRC LMB for plasmids containing several Cep proteins. The authors also thank J. Howe from the MRC LMB Light Microscopy for help with STORM, F. Begum from the MRC LMB Mass-Spectrometry for performing the BioID mass-spec runs and S. Zhang and S. Yatskevich for providing components and purified APC/C for in vitro ubiquitination assays. This work was supported by the Medical Research Council (MC_UP_1201/6) and a Cancer Research UK grant (C576/A14109) to D. Barford.

## Author Contributions

TT performed and analysed all experiments. JY purified proteins. TT and DB planned experiments and wrote the manuscript.

## Data availability

All data related to this study is presented within the manuscript and the supporting files. Cell lines that were generated for and used in this study are available upon request from the authors.

---

**Supplementary Figure 1.**
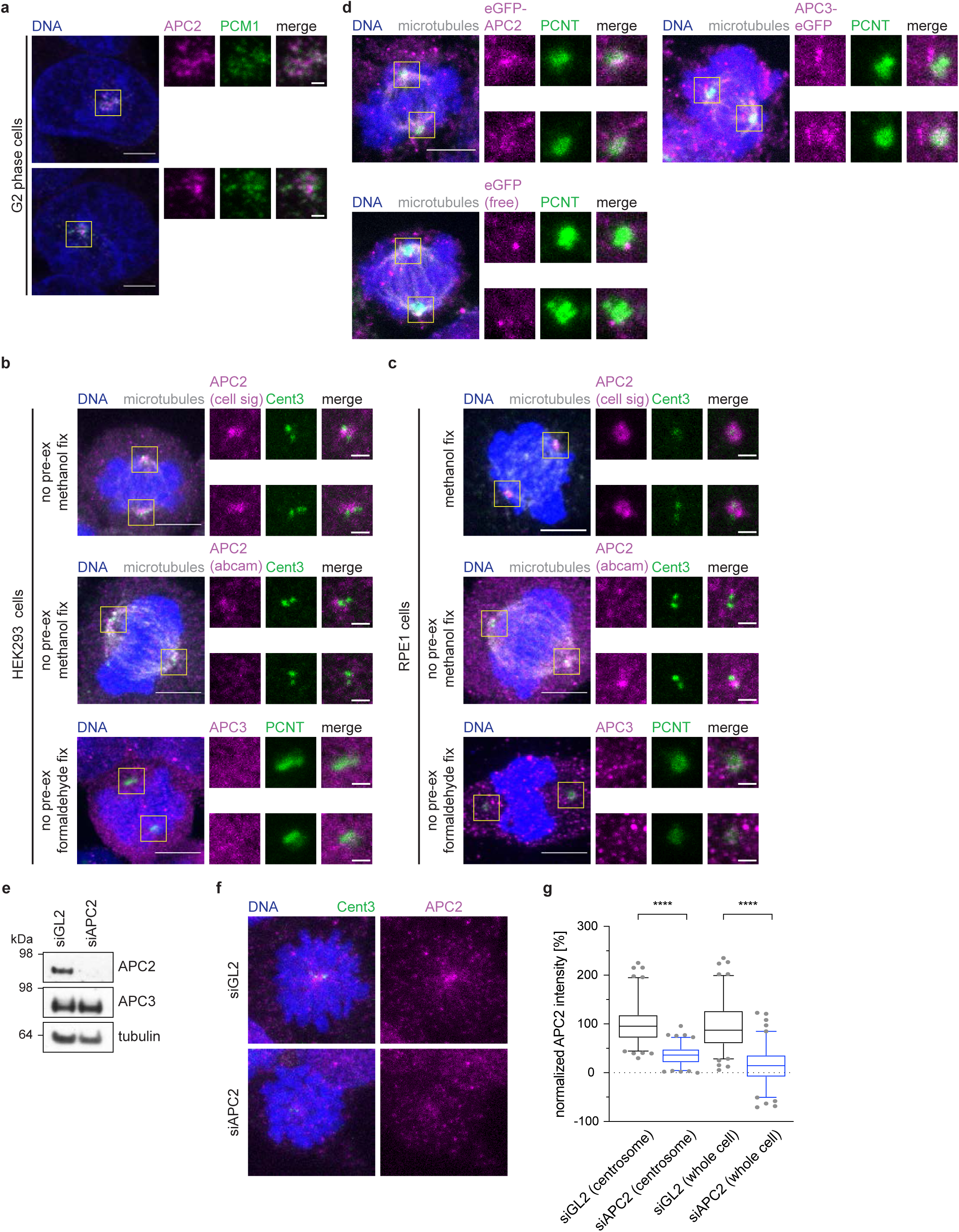
The APC/C localizes to centriolar satellites in interphase. **a**, HEK293 cells were release from a thymidine arrest for 4 hours, fixed with ice colde methanol and stained with the indicated antibodies. **b**, Mitotic HEK293 cells were not pre-extracted, fixed in ice cold methanol (top two rows) or formaldehyde (bottom) and stained with the indicated antibodies. **c**, RPE1 cells were either pre-extracted and fixed in ice-cold methanol (top row), not pre-extracted and fixed in ice-cold methanol (middle row) or not pre-extracted and fixed with formaldehyde (bottom) and subsequently stained with the indicated antibodies. **d**, HEK293 Flp In TRex cells harbouring the indicated eGFP-tagged transgenes were pre-extracted, fixed using formaldehyde, and stained with against the indicated proteins. **e**, Immunoblot from whole cell lysated of cells treated with control siRNA (siGL2) or siRNA against APC2 (siAPC2). **f**, Cells treated as in **e** were fixed in formaldehyde (no pre-extraction) and stained with the indicated antibodies. **g**, quantificationfrom cells shown in **f**. The measurment was either performed around the centrosome as defined by a 2 μm circle around the centrin3 signal or in the whole cell. All scale bars are 5 μm in the overviews and 1 μm in the insets.

**Supplementary Figure 2.**
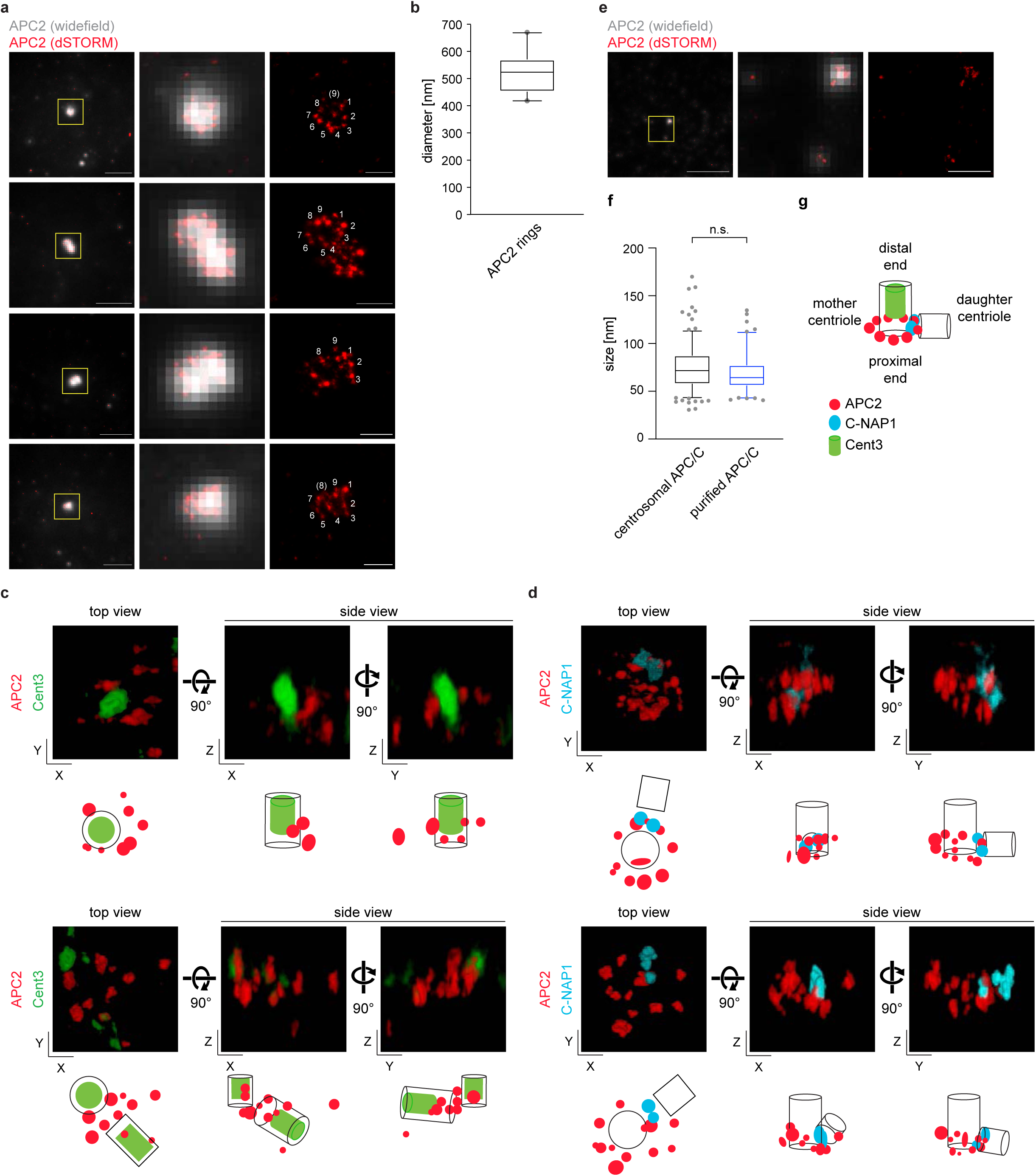
dSTORM imaging of the APC/C at mitotic centrosomes. **a**, HEK293 cells were arrested in prophase using taxol, pre-extracted and fixed with formaldehyde. APC2 was stained and dSTORM imaging was performed. The merged image shows the diffraction limited widefield image and the reconstructed dSTORM image. The numbers on the dSTORM image indicate the number of APC/C complexes that can be counted. Scale bars are 5 μm in the overview and 0.5 μm in the inset. **b**, The average diameter of the APC/C rings shown in **a** was measured in at least two directions per centrosome. N = 20. **c, d**, Cells were treated as described in **a**, but additionally stained against the indicated proteins. Two-color 3D STORM was performed using a cylindrical lens and by imaging both channels sequentially. Three different views are shown. The illustrations below the images indicate the possible orientation of the centrioles. See also Supplementary movies 1 - 4. **e**, Purified APC/C was fixed with formaldehyde. APC2 was stained and dSTORM imaging was performed. The merged image shows the diffraction limited widefield image and the reconstructed dSTORM image. Scale bars are 5 μm in the overview and 0.5 μm in the inset. **f**, The average diameter of the signal dots measured after STORM imaging of centrosomal APC/C or purified APC/C. N = 212 (centrosomal APC/C), 101 (purified APC/C). **g**, A model of the APC/C localization around the centrosome. The APC/C localizes within the PCM towards the proximal end of the centriole.

**Supplementary Figure 3.**
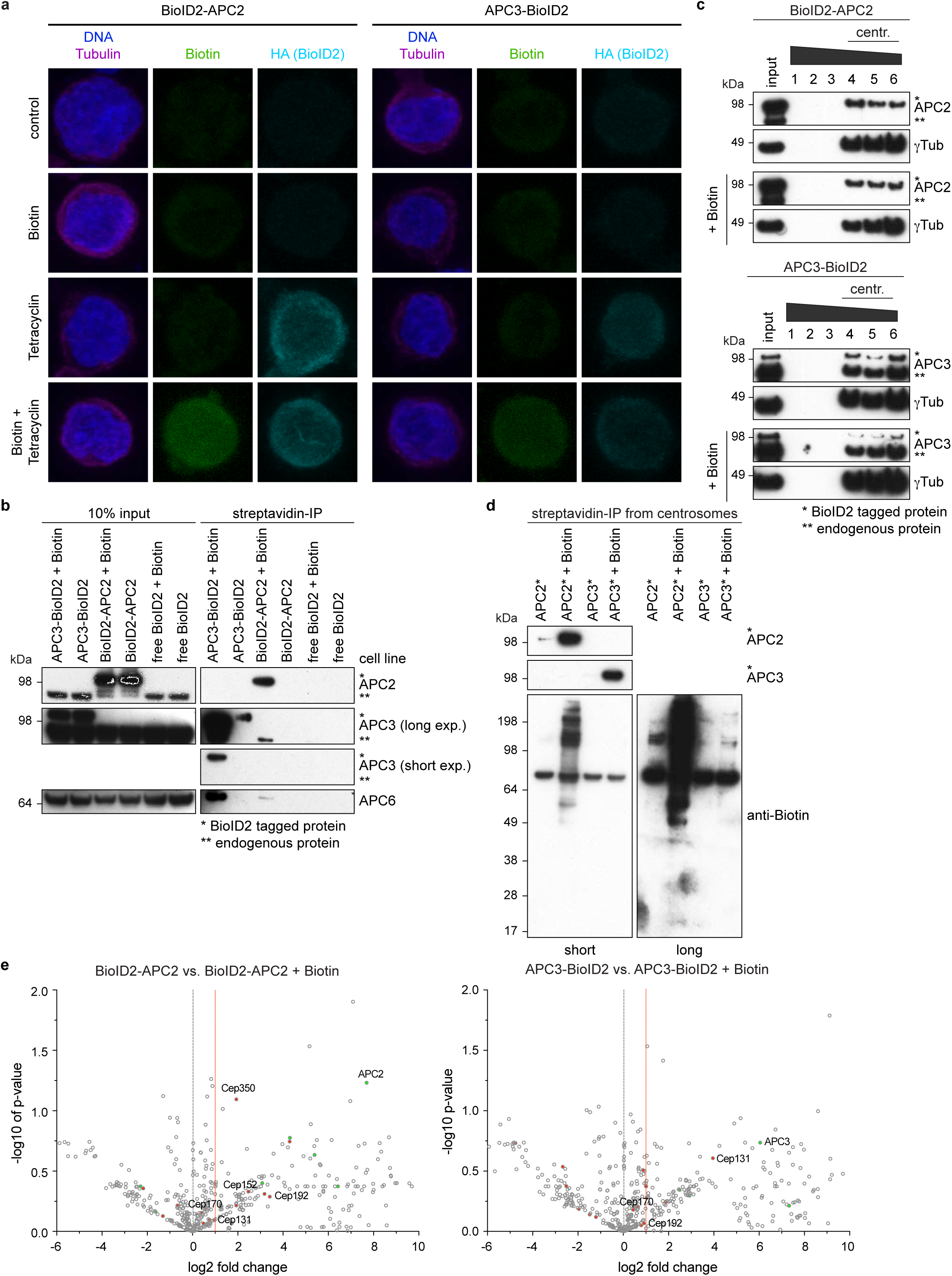
The APC/C is in close proximity to several other centrosomal proteins. **a**, HEK293 FlpIn T-Rex cells with stable integration of the indicated constructs were treated with the indicated compounds. **b**, HEK293 FlpIn T-Rex cells were treated with tetracycline to induce expression of the stably integrated BioID2 constructs and with or without biotin for proximity labelling. Whole cell lysates were subjected to streptavidin pulldown to enrich for biotinylated proteins. The samples were immunoblotted against the indicated proteins. **c**, Cells were treated as in **b**, but additionally blocked in mitosis by nocodazole, and mitotic centrosomes were purified via a sucrose gradient. The first elution fractions were immunoblotted against γTubulin as a centrosome marker and the APC/C. **d**, The centrosome containing fractions from **c** were pooled and subsequently used for a streptavidin pulldown to enrich specifically for biotinylated centrosomal proteins. **e**, The streptavidin pulldown from **d** was subjected to mass-spectrometry and the data were evaluated by label free analysis. The plots indicate centrosomal proteins with red dots and known APC/C interacting proteins with green dots. Proteins that were chosen for further analysis are indicated the graphs. See Supplementary Table 1 for the full mass-spec dataset.

**Supplementary Figure 4.**
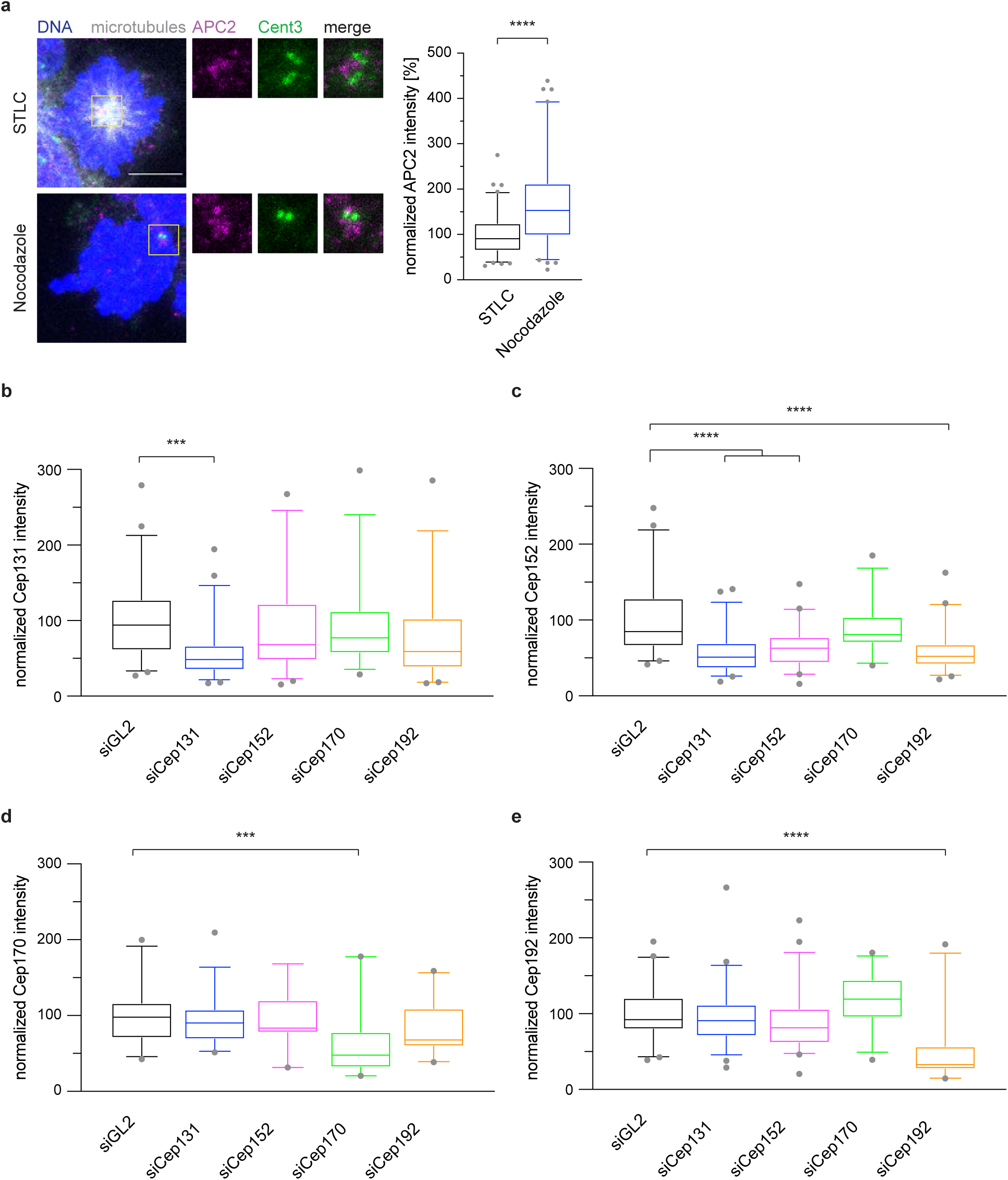
Depletion and cross-staining of centrosomal proteins identified by BioID2 mass-spectrometry. **a**, HEK293 cells were synchronized in mitosis using either STLC or Nocodazole, fixed and stained with the indicated antibodies (left). The fluorescence intensity of APC2 around the centrosome was anlyzed (right). **b, c, d, e**, HEK293 cells were depleted of the indicated centrosomal proteins using siRNA and arrested in mitosis by STLC. Cells were stained with the indicated antibodies and the centrosomal signal of each protein was quantified. The intensity was normalized against the siGL2 control. ****P < 0.0001 and ***P < 0.001 using a simple one-way ANOVA test with Dunnett’s multiple comparison. **a**, N = 48 (siGL2), 57 (siCep131), 49 (siCep152), 37 (siCep70), 39 (siCep192). **b**, N = 51 (siGL2), 50 (siCep131), 40 (siCep152), 29 (siCep70), 50 (siCep192). **c**, N = 29 (siGL2), 33 (siCep131), 19 (siCep152), 29 (siCep70), 22 (siCep192). **d**, N = 51 (siGL2), 46 (siCep131), 44 (siCep152), 33 (siCep70), 38 (siCep192).

**Supplementary Figure 5.**
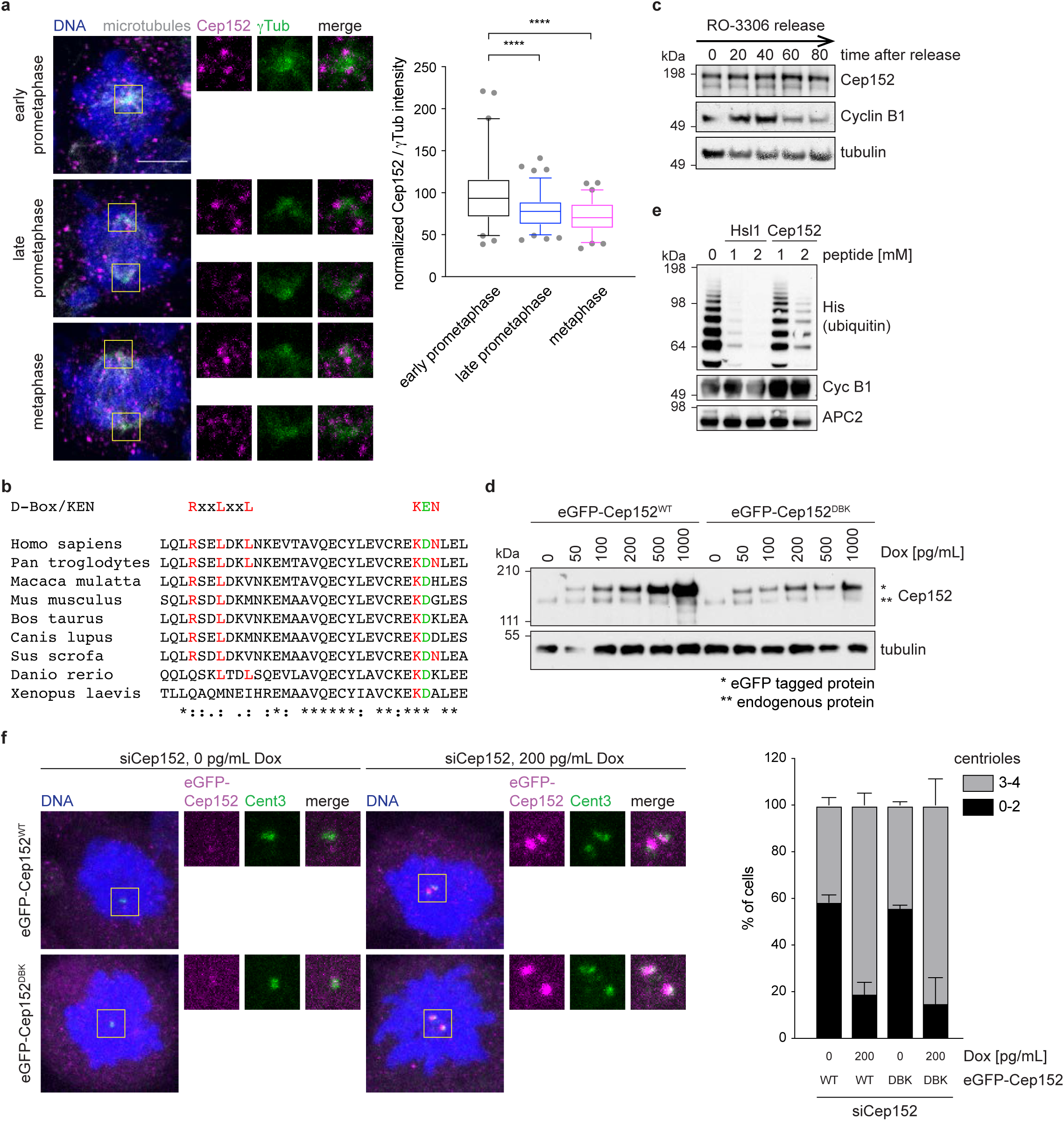
Cep152 protein levels at the centrosome decrease during mitosis. **a**, HEK293 cells were treated according to the time line in **Figure 1e** and stained against the indicated proteins. The fluorescence intensity of Cep152 at the centrosome was measured and normalized against the intensity in prophase. Scale bars are 5 μm in the overview and 1 μm in the inset. N = 72 (prophase), 92 (pro-metaphase), 68 (metaphase). ****P < 0.0001 using a simple one-way ANOVA test with Dunnett’s multiple comparison. **b**, Multiple sequence alignment of the Cep152 potential D box and KEN box. **c**, Cells were treated according to the time line in **Figure 1e** and whole cell lysates were prepared at the indicated time points. The lysates were probed by immunoblot against the indicated proteins. **d**, HEK293 Flp-In T-Rex cells carrying the indicated eGFP-Cep152 variants were treated with different concentrations of doxycycline for 24 h to induce the expression of the transgenes. Whole cell lysates were prepared and probed by immunoblot against the indicated proteins. Note that eGFP-Cep152 runs higher as the endogenous protein. For all other experiments 200 pg/mL doxycycline was used. **e**, Cyclin B1 was ubiquitinated in vitro using purified APC/C in the presence of increasing concentrations of Hsl1 or Cep152 peptide, respectively. **f**, HEK293 Flp-In T-Rex cells carrying the indicated eGFP-Cep152 variants were depleted of endogenous Cep152 by siRNA in the presence or absence of Doxycycline to induce the expression of the transgene according to the time line shown in **Figure 2a**. Cells were fixed with ice-cold methanol and stained with the indicated antibodies for immunofluorescence (left). The number of centrin-3 dots was counted and cells were classified as shown on the right. Depicted is the mean +/-s.d..

**Supplementray Figure 6.**
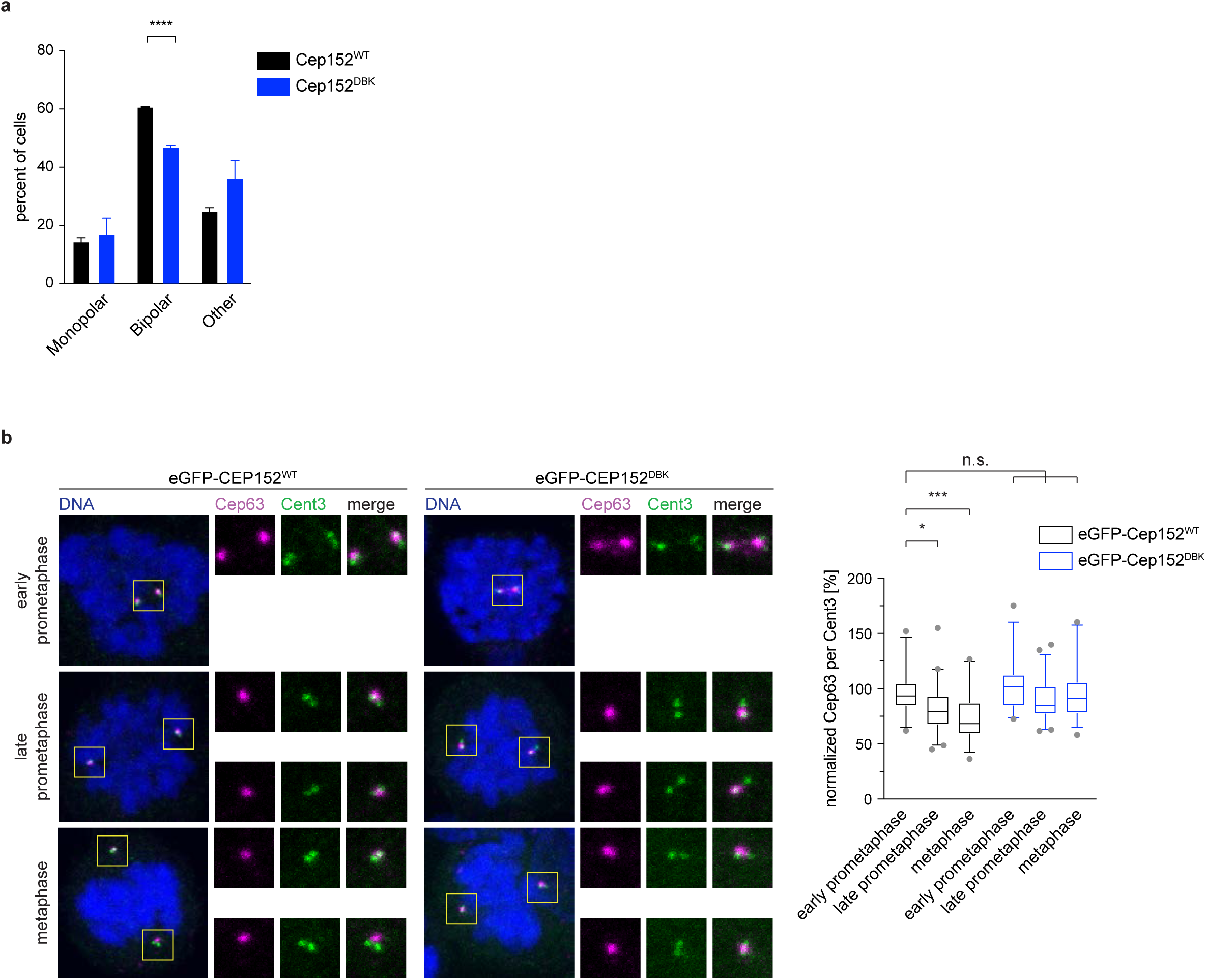
Reduced microtubule nucleation in the presence of stabilised Cep152. **a**, Cells shown in **Figure 4c** were analyzed for their spindle morphology after cold treatment for 45 min at 4 °C and recovery at 37 °C for 30 minutes. **b**, HEK293 FlpIn T-Rex cell lines expressing eGFP-Cep152 WT or DBK mutant protein were treated as shown in **Figure 1e** and stained with the indicated proteins (left). The fluorescence intensity of Cep63 was measured around the centrosome and normalized against prophase of the eGFP-Cep152^WT^ cell line (right). N = 32 (prophase, WT), 56 (pro-meta-phase, WT), 27 (metaphase, WT), 32 (prophase, DBK), 54 (pro-metaphase, DBK), 33 (metaphase, DBK). n.s. = not significant.*P < 0.05, *** P < 0.001, using a simple one-way ANOVA test with Dunnett’s multiple comparison.

## References

Acquaviva, C., Herzog, F., Kraft, C., and Pines, J. (2004). The anaphase promoting complex/cyclosome is recruited to centromeres by the spindle assembly checkpoint. Nature cell biology 6, 892–898.

Alfieri, C., Zhang, S., and Barford, D. (2017). Visualizing the complex functions and mechanisms of the anaphase promoting complex/cyclosome (APC/C). Open Biol 7.

Ban, K.H., Torres, J.Z., Miller, J.J., Mikhailov, A., Nachury, M.V., Tung, J.J., Rieder, C.L., and Jackson, P.K. (2007). The END network couples spindle pole assembly to inhibition of the anaphase-promoting complex/cyclosome in early mitosis. Developmental cell 13, 29–42.

Barenz, F., Kschonsak, Y.T., Meyer, A., Jafarpour, A., Lorenz, H., and Hoffmann, I. (2018). Ccdc61 controls centrosomal localization of Cep170 and is required for spindle assembly and symmetry. Molecular biology of the cell, mbcE18020115.

Betzig, E., Patterson, G.H., Sougrat, R., Lindwasser, O.W., Olenych, S., Bonifacino, J.S., Davidson, M.W., Lippincott-Schwartz, J., and Hess, H.F. (2006). Imaging intracellular fluorescent proteins at nanometer resolution. Science 313, 1642–1645.

Blachon, S., Gopalakrishnan, J., Omori, Y., Polyanovsky, A., Church, A., Nicastro, D., Malicki, J., and Avidor-Reiss, T. (2008). Drosophila asterless and vertebrate Cep152 Are orthologs essential for centriole duplication. Genetics 180, 2081–2094.

Clute, P., and Pines, J. (1999). Temporal and spatial control of cyclin B1 destruction in metaphase. Nature cell biology 1, 82–87.

Engelender, S., Sharp, A.H., Colomer, V., Tokito, M.K., Lanahan, A., Worley, P., Holzbaur, E.L., and Ross, C.A. (1997). Huntingtin-associated protein 1 (HAP1) interacts with the p150Glued subunit of dynactin. Human molecular genetics 6, 2205–2212.

Faragher, A.J., and Fry, A.M. (2003). Nek2A kinase stimulates centrosome disjunction and is required for formation of bipolar mitotic spindles. Molecular biology of the cell 14, 2876–2889.

Fong, C.S., Kim, M., Yang, T.T., Liao, J.C., and Tsou, M.F. (2014). SAS-6 assembly templated by the lumen of cartwheel-less centrioles precedes centriole duplication. Developmental cell 30, 238–245.

Fry, A.M., Meraldi, P., and Nigg, E.A. (1998). A centrosomal function for the human Nek2 protein kinase, a member of the NIMA family of cell cycle regulators. The EMBO journal 17, 470–481.

Gheiratmand, L., Coyaud, E., Gupta, G.D., Laurent, E.M., Hasegan, M., Prosser, S.L., Goncalves, J., Raught, B., and Pelletier, L. (2019). Spatial and proteomic profiling reveals centrosome-independent features of centriolar satellites. The EMBO journal 38, e101109.

Gomez-Ferreria, M.A., Rath, U., Buster, D.W., Chanda, S.K., Caldwell, J.S., Rines, D.R., and Sharp, D.J. (2007). Human Cep192 is required for mitotic centrosome and spindle assembly. Current biology : CB 17, 1960–1966.

Gonczy, P. (2012). Towards a molecular architecture of centriole assembly. Nature reviews Molecular cell biology 13, 425–435.

Gould, R.R., and Borisy, G.G. (1977). The pericentriolar material in Chinese hamster ovary cells nucleates microtubule formation. The Journal of cell biology 73, 601–615.

Guarguaglini, G., Duncan, P.I., Stierhof, Y.D., Holmstrom, T., Duensing, S., and Nigg, E.A. (2005). The forkhead-associated domain protein Cep170 interacts with Polo-like kinase 1 and serves as a marker for mature centrioles. Molecular biology of the cell 16, 1095–1107.

Gupta, G.D., Coyaud, E., Goncalves, J., Mojarad, B.A., Liu, Y., Wu, Q., Gheiratmand, L., Comartin, D., Tkach, J.M., Cheung, S.W., et al. (2015). A Dynamic Protein Interaction Landscape of the Human Centrosome-Cilium Interface. Cell 163, 1484–1499.

Gupta, G.D., and Pelletier, L. (2017). Centrosome Biology: Polymer-Based Centrosome Maturation. Current biology : CB 27, R836–R839.

Hori, A., and Toda, T. (2017). Regulation of centriolar satellite integrity and its physiology. Cell Mol Life Sci 74, 213–229.

Huang, J., and Raff, J.W. (1999). The disappearance of cyclin B at the end of mitosis is regulated spatially in Drosophila cells. The EMBO journal 18, 2184–2195.

Huang, T.T., Wuerzberger-Davis, S.M., Wu, Z.H., and Miyamoto, S. (2003). Sequential modification of NEMO/IKKgamma by SUMO-1 and ubiquitin mediates NF-kappaB activation by genotoxic stress. Cell 115, 565–576.

Izumi, H., Matsumoto, Y., Ikeuchi, T., Saya, H., Kajii, T., and Matsuura, S. (2009). BubR1 localizes to centrosomes and suppresses centrosome amplification via regulating Plk1 activity in interphase cells. Oncogene 28, 2806–2820.

Jehl, P., Manguy, J., Shields, D.C., Higgins, D.G., and Davey, N.E. (2016). ProViz-a web-based visualization tool to investigate the functional and evolutionary features of protein sequences. Nucleic Acids Res 44, W11–15.

Kallio, M.J., Beardmore, V.A., Weinstein, J., and Gorbsky, G.J. (2002). Rapid microtubule-independent dynamics of Cdc20 at kinetochores and centrosomes in mammalian cells. The Journal of cell biology 158, 841–847.

Kim, D.I., Jensen, S.C., Noble, K.A., Kc, B., Roux, K.H., Motamedchaboki, K., and Roux, K.J. (2016). An improved smaller biotin ligase for BioID proximity labeling. Molecular biology of the cell 27, 1188–1196.

Kim, J., Kim, J., and Rhee, K. (2019). PCNT is critical for the association and conversion of centrioles to centrosomes during mitosis. Journal of cell science 132.

Kim, S., and Rhee, K. (2014). Importance of the CEP215-pericentrin interaction for centrosome maturation during mitosis. PloS one 9, e87016.

Kim, T.S., Park, J.E., Shukla, A., Choi, S., Murugan, R.N., Lee, J.H., Ahn, M., Rhee, K., Bang, J.K., Kim, B.Y., et al. (2013). Hierarchical recruitment of Plk4 and regulation of centriole biogenesis by two centrosomal scaffolds, Cep192 and Cep152. Proceedings of the National Academy of Sciences of the United States of America 110, E4849–4857.

Kodani, A., Yu, T.W., Johnson, J.R., Jayaraman, D., Johnson, T.L., Al-Gazali, L., Sztriha, L., Partlow, J.N., Kim, H., Krup, A.L., et al. (2015). Centriolar satellites assemble centrosomal microcephaly proteins to recruit CDK2 and promote centriole duplication. eLife 4.

Kruse, J.-P., and Gu, W. (2009). MSL2 Promotes Mdm2-independent Cytoplasmic Localization of p53. Journal of Biological Chemistry 284, 3250–3263.

Kubo, A., Sasaki, H., Yuba-Kubo, A., Tsukita, S., and Shiina, N. (1999). Centriolar satellites: molecular characterization, ATP-dependent movement toward centrioles and possible involvement in ciliogenesis. The Journal of cell biology 147, 969–980.

Kubo, A., and Tsukita, S. (2003). Non-membranous granular organelle consisting of PCM-1: subcellular distribution and cell-cycle-dependent assembly/disassembly. Journal of cell science 116, 919–928.

Lawo, S., Hasegan, M., Gupta, G.D., and Pelletier, L. (2012). Subdiffraction imaging of centrosomes reveals higher-order organizational features of pericentriolar material. Nature cell biology 14, 1148–1158.

Lopes, C.A., Prosser, S.L., Romio, L., Hirst, R.A., O’Callaghan, C., Woolf, A.S., and Fry, A.M. (2011). Centriolar satellites are assembly points for proteins implicated in human ciliopathies, including oral-facial-digital syndrome 1. Journal of cell science 124, 600–612.

Lukinavicius, G., Lavogina, D., Orpinell, M., Umezawa, K., Reymond, L., Garin, N., Gonczy, P., and Johnsson, K. (2013). Selective chemical crosslinking reveals a Cep57-Cep63-Cep152 centrosomal complex. Current biology : CB 23, 265–270.

Mazo, G., Soplop, N., Wang, W.J., Uryu, K., and Tsou, M.F. (2016). Spatial Control of Primary Ciliogenesis by Subdistal Appendages Alters Sensation-Associated Properties of Cilia. Developmental cell 39, 424–437.

Meghini, F., Martins, T., Tait, X., Fujimitsu, K., Yamano, H., Glover, D.M., and Kimata, Y. (2016). Targeting of Fzr/Cdh1 for timely activation of the APC/C at the centrosome during mitotic exit. Nature communications 7, 12607.

Melan, M.A., and Sluder, G. (1992). Redistribution and differential extraction of soluble proteins in permeabilized cultured cells. Implications for immunofluorescence microscopy. Journal of cell science 101 (Pt 4), 731–743.

Merdes, A., Heald, R., Samejima, K., Earnshaw, W.C., and Cleveland, D.W. (2000). Formation of spindle poles by dynein/dynactin-dependent transport of NuMA. The Journal of cell biology 149, 851–862.

Merdes, A., Ramyar, K., Vechio, J.D., and Cleveland, D.W. (1996). A complex of NuMA and cytoplasmic dynein is essential for mitotic spindle assembly. Cell 87, 447–458.

Nigg, E.A., and Holland, A.J. (2018). Once and only once: mechanisms of centriole duplication and their deregulation in disease. Nature reviews Molecular cell biology.

Paoletti, A., Moudjou, M., Paintrand, M., Salisbury, J.L., and Bornens, M. (1996). Most of centrin in animal cells is not centrosome-associated and centrosomal centrin is confined to the distal lumen of centrioles. Journal of cell science 109 (Pt 13), 3089–3102.

Prosser, S.L., and Pelletier, L. (2020). Centriolar satellite biogenesis and function in vertebrate cells. Journal of cell science 133.

Quarantotti, V., Chen, J.X., Tischer, J., Gonzalez Tejedo, C., Papachristou, E.K., D’Santos, C.S., Kilmartin, J.V., Miller, M.L., and Gergely, F. (2019). Centriolar satellites are acentriolar assemblies of centrosomal proteins. The EMBO journal 38, e101082.

Raff, J.W., Jeffers, K., and Huang, J.Y. (2002). The roles of Fzy/Cdc20 and Fzr/Cdh1 in regulating the destruction of cyclin B in space and time. The Journal of cell biology 157, 1139–1149.

Rust, M.J., Bates, M., and Zhuang, X. (2006). Sub-diffraction-limit imaging by stochastic optical reconstruction microscopy (STORM). Nat Methods 3, 793–795.

Sivakumar, S., Daum, J.R., Tipton, A.R., Rankin, S., and Gorbsky, G.J. (2014). The spindle and kinetochore-associated (Ska) complex enhances binding of the anaphase-promoting complex/cyclosome (APC/C) to chromosomes and promotes mitotic exit. Molecular biology of the cell 25, 594–605.

Sonnen, K.F., Gabryjonczyk, A.M., Anselm, E., Stierhof, Y.D., and Nigg, E.A. (2013). Human Cep192 and Cep152 cooperate in Plk4 recruitment and centriole duplication. Journal of cell science 126, 3223–3233.

Sonnen, K.F., Schermelleh, L., Leonhardt, H., and Nigg, E.A. (2012). 3D-structured illumination microscopy provides novel insight into architecture of human centrosomes. Biol Open 1, 965–976.

Stearns, T. (2001). Centrosome duplication. a centriolar pas de deux. Cell 105, 417–420.

Thawani, A., Kadzik, R.S., and Petry, S. (2018). XMAP215 is a microtubule nucleation factor that functions synergistically with the gamma-tubulin ring complex. Nature cell biology 20, 575–585.

Tugendreich, S., Tomkiel, J., Earnshaw, W., and Hieter, P. (1995). CDC27Hs colocalizes with CDC16Hs to the centrosome and mitotic spindle and is essential for the metaphase to anaphase transition. Cell 81, 261–268.

Varmark, H. (2004). Functional role of centrosomes in spindle assembly and organization. Journal of cellular biochemistry 91, 904–914.

Watanabe, K., Takao, D., Ito, K.K., Takahashi, M., and Kitagawa, D. (2019). The Cep57-pericentrin module organizes PCM expansion and centriole engagement. Nature communications 10, 931.

Watson, E.R., Brown, N.G., Peters, J.M., Stark, H., and Schulman, B.A. (2019). Posing the APC/C E3 Ubiquitin Ligase to Orchestrate Cell Division. Trends Cell Biol 29, 117–134.

Wei, Z., Kim, T.S., Ahn, J.I., Meng, L., Chen, Y., Ryu, E.K., Ku, B., Zhou, M., Kim, S.J., Bang, J.K., et al. (2020). Requirement of the Cep57-Cep63 Interaction for Proper Cep152 Recruitment and Centriole Duplication. Molecular and cellular biology 40.

Wieczorek, M., Urnavicius, L., Ti, S.C., Molloy, K.R., Chait, B.T., and Kapoor, T.M. (2020). Asymmetric Molecular Architecture of the Human gamma-Tubulin Ring Complex. Cell 180, 165–175 e116.

Woodruff, J.B., Ferreira Gomes, B., Widlund, P.O., Mahamid, J., Honigmann, A., and Hyman, A.A. (2017). The Centrosome Is a Selective Condensate that Nucleates Microtubules by Concentrating Tubulin. Cell 169, 1066–1077 e1010.

Woodruff, J.B., Wueseke, O., and Hyman, A.A. (2014). Pericentriolar material structure and dynamics. Philosophical transactions of the Royal Society of London Series B, Biological sciences 369.

Wu, Q., He, R., Zhou, H., Yu, A.C., Zhang, B., Teng, J., and Chen, J. (2012). Cep57, a NEDD1-binding pericentriolar material component, is essential for spindle pole integrity. Cell Res 22, 1390–1401.

Yan, X., Habedanck, R., and Nigg, E.A. (2006). A complex of two centrosomal proteins, CAP350 and FOP, cooperates with EB1 in microtubule anchoring. Molecular biology of the cell 17, 634–644.

Yao, F., Zhou, Z., Kim, J., Hang, Q., Xiao, Z., Ton, B.N., Chang, L., Liu, N., Zeng, L., Wang, W., et al. (2018). SKP2- and OTUD1-regulated non-proteolytic ubiquitination of YAP promotes YAP nuclear localization and activity. Nature communications 9.

Zhang, S., Chang, L., Alfieri, C., Zhang, Z., Yang, J., Maslen, S., Skehel, M., and Barford, D. (2016). Molecular mechanism of APC/C activation by mitotic phosphorylation. Nature 533, 260–264.

Zhang, S., Tischer, T., and Barford, D. (2019). Cyclin A2 degradation during the spindle assembly checkpoint requires multiple binding modes to the APC/C. Nature communications 10, 3863.

Zhao, H., Yang, S., Chen, Q., Duan, X., Li, G., Huang, Q., Zhu, X., and Yan, X. (2020). Cep57 and Cep57l1 function redundantly to recruit the Cep63-Cep152 complex for centriole biogenesis. Journal of cell science 133.

Zhou, Y., Ching, Y.P., Chun, A.C., and Jin, D.Y. (2003). Nuclear localization of the cell cycle regulator CDH1 and its regulation by phosphorylation. The Journal of biological chemistry 278, 12530–12536.

Zybailov, B., Mosley, A.L., Sardiu, M.E., Coleman, M.K., Florens, L., and Washburn, M.P. (2006). Statistical analysis of membrane proteome expression changes in Saccharomyces cerevisiae. J Proteome Res 5, 2339–2347.

